# Sleep pressure modulates single-neuron synapse dynamics in zebrafish

**DOI:** 10.1101/2023.08.30.555615

**Authors:** Anya Suppermpool, Declan G. Lyons, Elizabeth Broom, Jason Rihel

## Abstract

Sleep is a nearly universal behaviour with unclear functions_1_. The Synaptic Homeostasis Hypothesis (SHY) proposes that sleep is required to renormalize the increases in synaptic number and strength that occur during wakefulness_2_. Some studies examining either large neuronal populations_3_ or small patches of dendrites_4_ have found evidence consistent with SHY, but whether sleep merely serves as a permissive state or actively promotes synaptic downregulation at the scale of whole neurons is unknown. Here, by repeatedly imaging all excitatory synapses on single neurons across sleep/wake states of zebrafish larvae, we show that synapses are gained during periods of wake (either spontaneous or forced) and lost during sleep in a neuron-subtype dependent manner. However, synapse loss is greatest during sleep associated with high sleep pressure following prolonged wakefulness and low in the latter half of the night. Conversely, sleep induced pharmacologically during periods of low sleep pressure is insufficient to trigger synapse loss unless adenosine levels are boosted while noradrenergic tone is inhibited. We conclude that sleep-dependent synapse loss is regulated by sleep pressure at the level of the single neuron and that not all sleep periods are equally capable of fulfilling the functions of synaptic homeostasis.

## Introduction

Although sleep is conserved across the animal kingdom^1^, the precise functions of sleep remain unclear. Since sleep deprivation leads to acute impairment of cognitive performance^5^, many theories posit that synaptic plasticity associated with learning and memory preferentially occurs during sleep^6^. For example, the Synaptic Homeostasis Hypothesis (SHY) proposes that synaptic potentiation during wakefulness results in an ultimately unsustainable increase in synaptic strength and number that must be renormalized during sleep through synaptic weakening and pruning^2,7,8^. Such sleep-dependent renormalization has been postulated to broadly affect most excitatory synapses throughout the brain^2^.

Many, but not all, experimental observations of brain-wide changes in synapses have been consistent with SHY. Globally, synaptic genes, proteins, and post-translational modifications are upregulated during waking and renormalize during sleep^9–12^. In both flies and mice, the number and size of excitatory synapses also increase after prolonged waking and decline during sleep^3,10,13^. Long term imaging of small segments of dendrites in young and adult mice have also observed sleep/wake-linked synapse dynamics^4,14,15^, and in zebrafish, axon terminals of wake-promoting hypocretin neurons are circadian-clock regulated to peak during the day^16^. However, other studies have observed no impact of sleep/wake states on synaptic strength and neuronal firing rates^17,18^, and some have observed synaptic strengthening during sleep^19–22^. Furthermore, distinct classes of synapse within the same neuronal population can be differentially regulated by sleep/wake states^23^, consistent with observations that synaptic plasticity can be regulated in a dendritic branch-specific manner^24^. Together, these observations paint a complex picture of how sleep sculpts synapse dynamics, raising fundamental questions about whether sleep-dependent synaptic homeostasis operates uniformly across neuronal types and on which scale (e.g., dendrite, neuron, circuit, or population) sleep acts to modulate synapses.

To examine the scope and selectivity of sleep-linked synaptic plasticity, it is vital to comprehensively track the synaptic changes of individual neurons through sleep/wake states. To that end, we used *in vivo* synaptic labelling tools in larval zebrafish to image the same neurons and their synapses repeatedly over long timescales, allowing us to map single-neuron synapse dynamics across sleep and wake states.

## Results

### Single neuron synapse dynamics across day:night cycles

To visualize excitatory synapses in single zebrafish neurons, we adapted an established Fibronectin intrabodies generated with mRNA display (FingR) based transgenic system that selectively binds to and labels PSD-95^25–27^, a major postsynaptic scaffold of excitatory synapses^28,29^ and a readout of synaptic strength^30,31^, to allow for simultaneous imaging of synapses and neuronal morphology (**Figure 1a**). Consistent with previous reports^25,27,32^, we confirmed this modified FingR(PSD95)+ system labels synapses with high fidelity by driving expression of *Tg(UAS:FingR(PSD95)-GFP-P2A-mKate2f)* in the spinal cord with a *Tg(mnx1:Gal4)* driver line and co-labelling with anti-MAGUK antibodies that recognize the PSD-95 protein family. Greater than >90% of FingR(PSD95)+ puncta associated with MAGUK, while 100% of neuronal MAGUK puncta were co-labelled with FingR(PSD95) (**Extended Data Figure 1a-e, h-i**). The signal intensities of co-labelled MAGUK and FingR(PSD95) synapses were positively correlated, indicating that signal intensity is a reliable readout of synaptic strength as reported (**Extended Data Figure 1f,g**)^26^.

**Figure 1:**
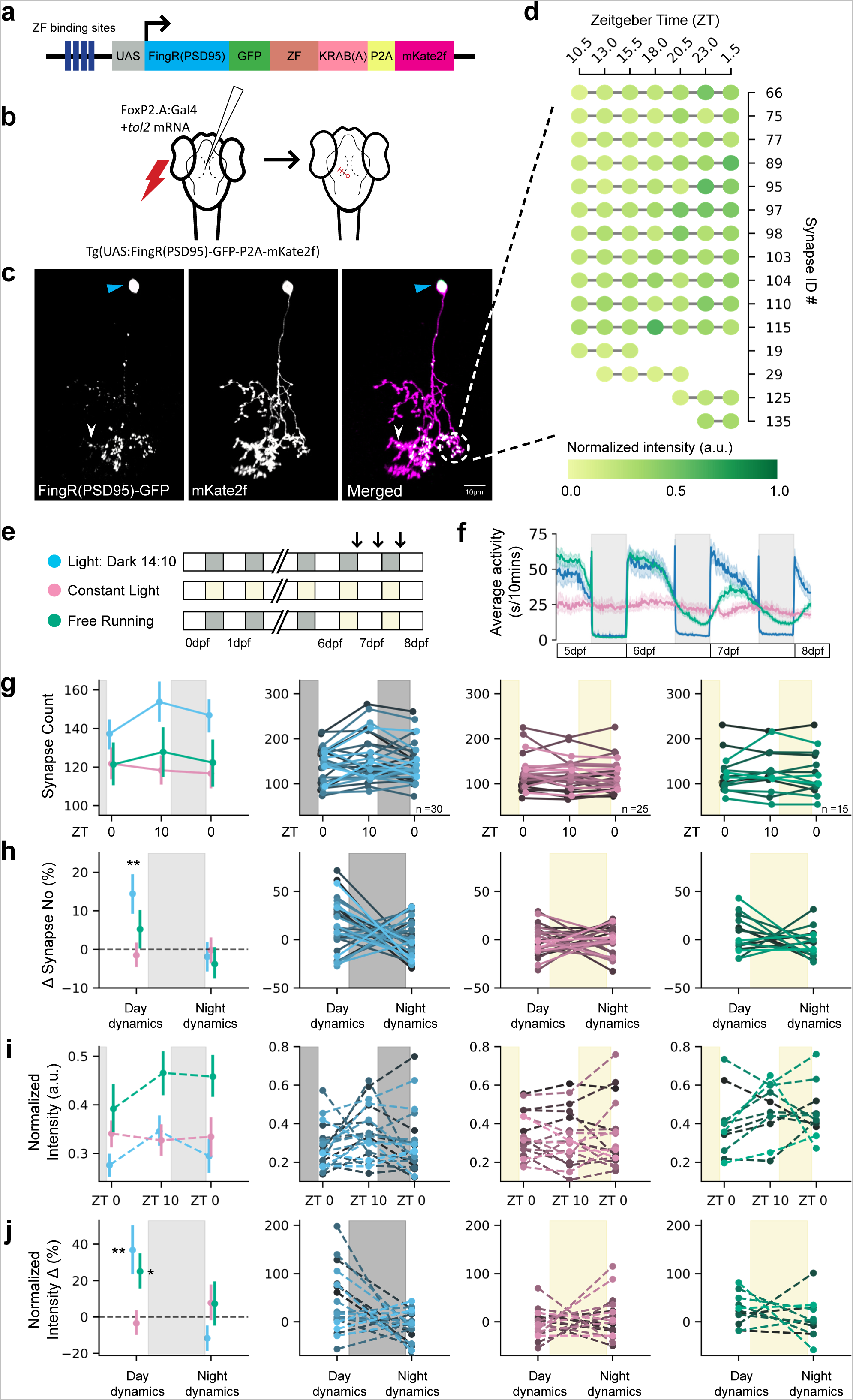
Single neuron synapse tracking across day:night cycles reveals diverse dynamics. **a**, Schematic of the UAS:FingR(PSD95) and membrane targeting mKate2f construct. The zinc finger (ZF) domain directs unbound FingR(PSD95) proteins to supress its own expression via an inhibitory KRAB(A) domain^25^. **b**, Schematic of the electroporation strategy to label synapses of FoxP2.A+ tectal neurons. Tol2 allows for genomic integration of the FoxP2.A:Gal4 plasmid, allowing for sparse, stable expression. See **Methods** for details. **c**, An example of a FoxP2.A:FingR(PSD95)+ neuron at 7 dpf, with synapses (white arrowhead, **left**), nucleus (blue arrowhead, **left**), and membrane (magenta, **middle**) co-labelled. **d**, Selection of overnight (between ZT10 to ZT1) time-lapse synapse tracking of the same neuron in **c**. Each row depicts a synapse, and the colour indicates the normalized GFP intensity of each synapse. See **Extended Data Figure 2a** for the complete map of overnight changes in all 138 synapses. **e**, Larvae were raised either on 14hr:10hr light:dark (LD) cycles (blue), on constant light (LL, pink), or switched from LD to LL at 6 dpf (‘free running’, FR, green). The arrows show the times of synapse imaging (see **Extended Data Table 2**). **f**, Average locomotor activity through multiple days and nights of larvae reared in normal LD (blue, n=75), ‘clock-breaking’ LL (pink, n=84), or FR (green, n=98) conditions, as depicted in **e**. **g**, The mean and 68% confidence interval (CI) of synapse counts at each timepoint in LD (blue), LL (pink), or FR (green) conditions (**left**). Synapse counts for each neuron are plotted as a single line (**right**). **h**, The percentage change (mean and 68% CI, **left**; each neuron, **right**) in synapse number calculated within each neuron across time (from **g**). There is a significant day:night difference (*P<0.05, repeated measures ANOVA) for LD dynamics, and LD cycling is different from LL conditions (**P<0.01, Mixed ANOVA with post-hoc pairwise t-test). **i**, The mean and 68% CI (**left**) of normalized synapse intensity. Each neuron is plotted separately on the **right**. **j**, The percentage change (mean and 68% CI, **left**; each neuron, **right**) in normalized synapse intensity calculated as in **h**. Day:night dynamics are significantly different in the LD (**P<0.01, repeated measures ANOVA) condition, and both FR and LD are significantly different from LL (FR-LL*P<0.05, LD-LL **P<0.01, Mixed ANOVA with pairwise correlation).

To test whether behavioural state modulates synapse strength and number at the single-neuron level, we focused on larval tectal neurons, which are accessible to imaging, have well-defined morphological and functional identities^33^, and have a stable window of synapse maturation from 7 to 9 days post fertilization (dpf)^34^. Tectal neurons also undergo spike-timing-dependent plasticity^35^ and receive a mixture of inputs that foster ‘competition’ among synapses^36,37^, a criterion envisaged by SHY^2^. To sparsely label tectal neurons, we co-electroporated a plasmid driving Gal4 off the *foxP2.A* promoter with *Tol2* mRNA into *Tg(UAS:FingR(PSD95)-GFP-P2A-mKate2f)* larvae at 3 dpf (**Figure 1b,c and Methods**)^38^. This method resulted in approximately 10% of larvae containing a single FoxP2.A:FingR(PSD95)+ neuron, allowing for repeated, long-term imaging of the synapse counts on and intensities of the same neuron in a continuously mounted preparation (**Figure 1c,d and Extended Data Figure 2**). After confirming the relative stability of tectal neuron synapse counts in the 6-9 dpf developmental window (**Extended Data Figure 2b-d**), we imaged each labelled neuron across a 14hr:10hr light:dark cycle at 7 dpf, collecting images just after lights on (Zeitgeber Time, ZT0, 7dpf), near the end of the day (ZT10), and after a night of sleep (ZT0, 8dpf) (**Figure 1e; see Extended Data Figure 3** for an example neuron with synapse changes tracked across two timepoints), leaving larvae to behave freely between imaging sessions. On average, tectal neuron synapse number significantly increased during the day from 137 to 153 synapses (+14.4%) and then decreased at night by -1.90% to 146 synapses (**Figure 1g**, **h**, blue). Similar day:night synapse dynamics were observed in separate experiments that imaged neurons over multiple days and nights (**Extended Data Figure 4a-e**), with no evidence for artefacts from repeated imaging (**Extended Data 4f-h**). Additionally, the average synapse GFP signal intensity significantly increased during the waking day phase (+36.8%) and decreased in the night sleep phase (-11.7%) (**Figure 1i-j**).

To test if these synaptic dynamics are influenced by the direct action of lighting conditions or are instead controlled by an internal circadian clock, we also tracked neurons under conditions of either constant light from fertilization, which prevents the formation of functional circadian clocks and leads to arrhythmic behaviour in zebrafish (‘clock break’)^39–41^, or constant light after entrainment, which maintains damped circadian behaviour (‘free running’) (**Figure 1f**)^42^. Under clock-break conditions, synapse dynamics in number and intensity were abolished and remained lower than larvae raised on light:dark cycles (**Figure 1g-j**, pink). Under free running conditions, synapse numbers continued to increase during the subjective day and decrease during the subjective night, albeit strongly damped (**Figure 1g**,**h**, green). The average synapse intensity was significantly elevated across all timepoints and showed a further significant increase in strength only during the subjective day, with no loss of intensity during the subjective night (**Figure 1i**,**j**, green). Collectively, these data show that, while light influences the baseline levels of synaptic strength (e.g. **Figure 1i**), changes in synapse counts are independent of lighting conditions but do require an intact circadian clock (to drive rhythmic sleep/wake behaviour, see below) (**Figure 1g**).

Moreover, although rhythmic day:night changes in synapses were detected in the average of all the single neurons, the tracking of individual neurons revealed that many cells have different, even opposing, synaptic dynamics (**Figure 1g-j**, right panels). We therefore sought to test whether these diverse patterns mapped onto distinct neuronal subtypes (i.e. cellular diversity) or might be due to variations in animal behaviour (i.e. individual sleep/wake histories).

### Tectal neuron subtypes have distinct synapse dynamics

To test whether distinct synapse day:night dynamics are associated with morphological subtypes of tectal neurons, we measured position, branching, length, and other parameters of FoxP2.A:FingR(PSD95)-GFP+ neurons that project only within the tectum at 7 dpf. Clustering analysis found four subtypes, consistent with previous work^33,43^ (**Figure 2a-c and Extended Data Figure 5a-c**). Tracking synapses across three light:dark cycles revealed that each neuronal subtype has, on average, different synapse dynamics (excluding the rarely observed Type 1 neurons). Specifically, dynamics consistent with SHY were only robustly observed in the densely bistratified Type 2 neurons, with an average increase of 14.3 synapses during the day and a reduction of 17.7 synapses at night, and weakly observed in Type 4 neurons (+8.5 during the day and -8.2 overnight; **Figure 2d-g and Extended Data Figure 5d-f**). In contrast, many Type 3 neurons consistently exhibited the opposite dynamic, with an average increase in synapse number at night and a slight decrease during the day (**Figure 2d-g**). These subtype-specific synapse dynamics cannot be explained by differences in larval sleep/wake behaviour, as sleep amount was the same regardless of which neuron sub-type was labelled in the larva (**Extended Data Figure 6b**).

**Figure 2:**
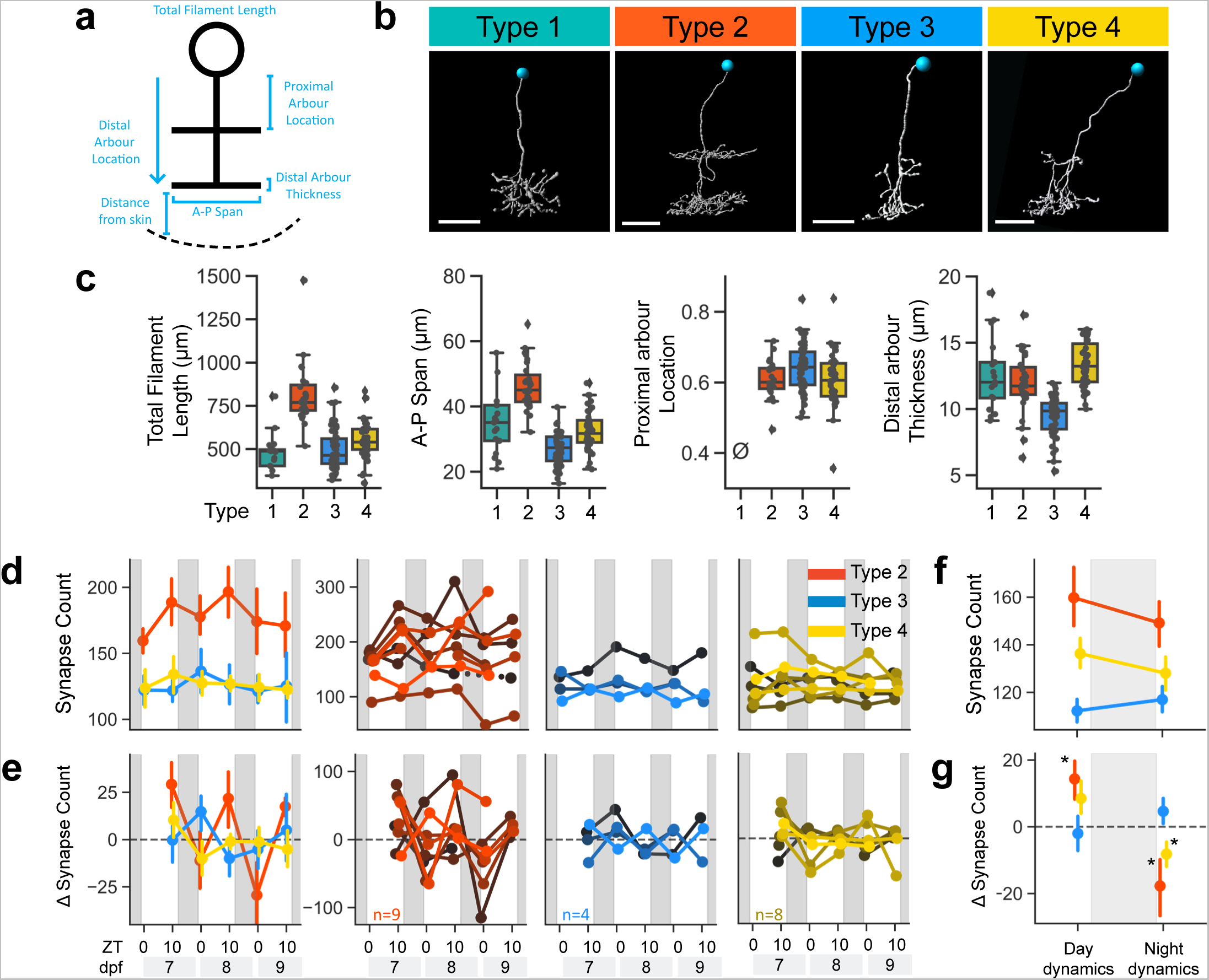
Subtype-specific synapse dynamics in FoxP2.A tectal neurons over three days. **a**, Morphological parameters used to characterize FoxP2.A tectal neurons. **b**, An example of each morphological subtype. Blue circles label nuclei. Scale bar, 10μm. **c**, Example of parameters used to distinguish the four subtypes. Boxes depict the median and interquartile range and the whiskers represent the distribution for each parameter. The slashed zero indicates the feature is absent. See **Extended Data Figure 5** for other parameters. **d-g**, Synapse counts across multiple LD cycles for FoxP2.A tectal neurons of different subtypes. **d**, Average synapse counts with 68% CI and **e**, average synapse number change of subtypes (**left**) and for each neuron (**right**). **f-g**, Average synapse counts and change, averaged across all days and nights for each subtype and larvae, and including additional neurons tracked over a single day (**Extended Data Figure 6**). Type 2 neurons (n=16) significantly gain synapses during the day, while Type 3 (n=14) and Type 4 neurons (n=15) were not different from zero (p=0.057 for Type4). At night Type 2 and Type 4 neurons significantly lose synapses, while Type 3 neurons were not different from zero (*P<0.05, directional one sample t-test).

Since Type 2 neurons have two prominent arborization fields, we asked whether day:night synapse dynamics are heterogenous across different dendritic segments of individual neurons. Analysing the synapse dynamics within each of four Type 2 dendritic segments revealed that only the proximal arbour, which receives local inputs from the tectum and long-range inputs from brain areas such as the hypothalamus^44^, displayed significantly robust average increases in synapse number during the day and reductions at night (**Extended Data Figure 7a-c**). In contrast, synapse number dynamics within the distal arbour, which receives the majority of its inputs from the retina^43^, were more diverse. No correlations could be detected among the different dendritic compartments within the same neuron (**Extended Data Figure 7d-e**), suggesting that time of day and sleep/wake states do not have uniform effects on synapse dynamics even within the same neuron.

### High sleep pressure facilitates sleep-dependent synapse loss

If single neuron synapse dynamics are regulated by sleep/wake states independently of the circadian clock, these dynamics should be altered by sleep deprivation (SD). We developed a gentle handling SD protocol in which zebrafish larvae are manually kept awake with a paintbrush for 4 hours at the beginning of the night (ZT14-18) and subsequently allowed to sleep (**Extended Data Video 1**). Sleep in larval zebrafish is defined as a period of inactivity lasting longer than one minute, as this is associated with an increased arousal threshold, homeostatic rebound, and other criteria of sleep^40,45^. After SD, the phase of the circadian clock machinery was unaffected, but larvae slept significantly more, with individual sleep bouts lasting longer, compared to non-deprived larvae (**Extended Data Figure 8a,b**), consistent with SD leading to increased sleep pressure^46–48^. Next, we visualized synapses of individual tectal neurons at 7 dpf immediately before (ZT13-14) and after (ZT18-20) SD, and again the following morning (ZT0-1) (**Figure 3a**, arrows). Between imaging sessions, we used video tracking to monitor larval sleep/wake behaviour (**Methods**). In control larvae, tectal neurons lost synapses overnight; however, this synapse loss was confined to the first part of the night (ZT14-18), with an average loss of 1.7 synapses/hr, compared to the last part of the night (ZT18-24), when synapse loss was undetectable (+0.2 synapses/hr) (**Figure 3b**, blue). In contrast, neurons gained an average of 2.8 synapses/hr during SD (**Figure 3b**, orange). During the post-SD recovery period, tectal neurons lost synapses at a rate of 2.2 synapses/hr (**Figure 3b and Extended Data Figure 8c).** As during normal sleep, FoxP2.A tectal neuron subtypes responded differently to SD, with Type 2, and even Type 3 neurons (which did not have SHY-concordant dynamics under baseline conditions), gaining synapses during SD and losing them during recovery sleep, whereas Type 4 neurons did not show any change (**Extended Data Figure 8d**). This suggests that SD biases synapses towards loss during subsequent sleep, even in neurons with different synapse dynamics in baseline conditions.

**Figure 3:**
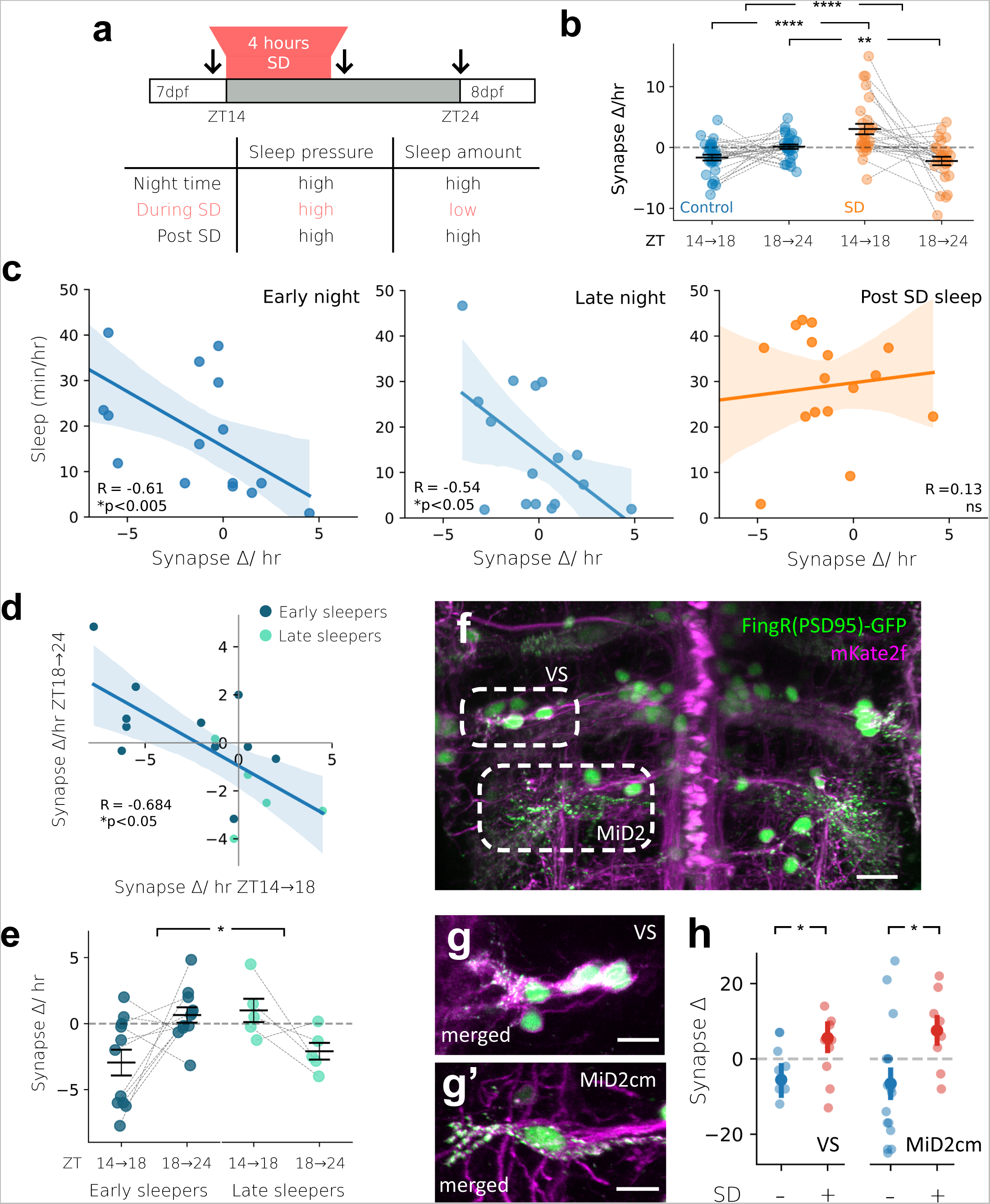
Synapse dynamics of neurons are modulated by sleep and sleep deprivation. **a**, The 4 hr gentle handling sleep deprivation paradigm (ZT14-18). Arrows indicate the imaging periods. Larvae were videotracked between imaging sessions. Gentle handling SD increases sleep pressure (i.e., high) while minimising sleep amount (i.e., low). **b**, Changes and ±SEM in synapse counts/hr for SD (orange, n=31) and control (blue, n=28) groups. **c**, For each neuron/larva, sleep time is plotted relative to the change in synapse counts/hr during either the early night (ZT14-18, **left**) or late night (ZT18-24, **middle**) for controls and after SD (ZT18-24, **right**). Rate of synapse change is negatively correlated with sleep time during both early and late night but not following SD. **d**, In control larvae, the change in synapse counts early in the night is negatively correlated with the synapse change in the late night. Early and late sleepers are defined as larvae that either sleep more in the first or second phase of the night, respectively. **e**, Synapse counts/hr for early-and late-night sleeping control larvae in the early (ZT14-18) and late (ZT18-24) phases of the night. The rare Type 1 neurons were excluded. **f-h**, The synapse dynamics of reticulospinal neurons are modulated by sleep and wake states. **f**, An example of reticulospinal neurons co-labelled by FingR(PSD95)-GFP (green, nuclei and synapses) and mKate2f (magenta, membrane). A labelled vestibulospinal (VS) and MiD2cm touch evoked neuron are indicated by dotted ovals. Scale bar, 15μm. **g-g’**, VS and MiD2cm neurons from a different larvae showing FingR(PSD95)+ synapses (green) co-localized to the cell membrane (magenta). Scale bar, 10μm. **h**, The change in synapse number (average and 68% CI) between ZT14 and ZT18 of vestibulospinal and MiD2cm neurons in control and SD larvae. Each dot represents the average across multiple neurons of the same type per larva. ****P<0.0001, **P<0.01, and *P<0.05 mixed ANOVA interaction and post-hoc pairwise t-test for **b** and **e;** *P<0.05, Mann-Whitney, one-tailed for **h**.

Since both SD and control larvae were at the same circadian phase, we conclude that sleep/wake states are the main driver of synapse dynamics in tectal neurons, and the effects of circadian clock disruption on synapses were primarily due to the loss of sleep rhythms (**Figure 1**). Consistent with this interpretation, the total time each larva spent asleep was significantly correlated with the rate of synapse change (**Figure 3c** and **Extended Data Figure 8g**). Only after SD, when sleep and synapse loss were high across most larvae-neuron pairs, was this correlation lost, which may indicate that either the machinery that supports sleep-dependent synapse loss can saturate or SD-induced rebound sleep is not fully equivalent to baseline sleep. The converse relationship was not observed: the rate of synapse gain during SD did not correlate with either the subsequent total sleep or the average sleep bout lengths of single larvae (**Extended Data Figure 8f**). Consistent with gentle handling SD, natural individual variation in sleep timing was predictive of the time period in which synapses were lost. ‘Early sleepers’ slept more in the first half of the night and lost synapses only during this period, while ‘late sleepers’ preferentially slept in the second half of the night and had a net loss of synapses only during the late night (**Figure 3d-e**; **Extended Data Figure 8e**). Finally, to test whether sleep-dependent synapse loss is generalizable to neurons that do not receive visual input, we confirmed that synapses of both vestibulospinal neurons that stabilize posture^49^ and MiD2cm reticulospinal neurons involved in fast escapes^50,51^ showed synapse gains during SD and synapse loss during sleep (**Figure 3f-h**).

Two explanations are consistent with the observed relationships between sleep and synapse dynamics: either sleep is a permissive state for synapse loss; or sleep pressure, which builds as a function of waking, drives synapse loss during subsequent sleep. Since sleep pressure and subsequent sleep amount at night are tightly linked under both baseline and SD conditions, we sought to disentangle their relative influences on synaptic change by using sleep-inducing drugs to force larvae to sleep during the day, when sleep pressure remains low (**Figure 4a-b**, **Extended Data Figure 9a**). Exposing larvae for 5 hrs during the day (ZT5-10) to either 30 µM melatonin, which in zebrafish is a natural hypnotic that acts downstream of the circadian clock to promote sleep^52^, or 30 µM clonidine, an α2-adrenergic receptor agonist that inhibits noradrenaline release and increases sleep in zebrafish^45,53^, significantly and strongly increased total sleep and the average length of sleep bouts mid-day (**Figure 4c**, **Extended Data Figure 10a-b**), with this drug-induced sleep remaining reversible by strong stimuli (**Extended Data Figure 9b-c and 11**). Forced daytime sleep altered the build-up of sleep pressure, leading to reduced and delayed sleep in the subsequent night (**Extended Data Figure 9c**). However, drug-induced sleep at a time of low sleep pressure was not sufficient to trigger synapse loss, with tectal neurons still gaining an average of 1.0-1.7 synapses/hr, which was not significantly different from the synapse gains in controls (**Figure 4d**). Similarly, artificially boosting adenosine signalling – one of the postulated molecular substrates of sleep pressure^54^ – by administering 45 µM 2-choloroadenosine increased sleep during the day but also led to net gains in tectal neuron synapses (+0.9 synapse/hr) (**Figure 4c** and **Extended Data Figure 10a-b**). Tectal neurons also gained synapses (+0.4synapse/hr) in larvae co-administered 2-chloroadenosine and melatonin, despite sleeping more than 35 min/hr (**Figure 4c**,**d**). In contrast, simultaneously boosting adenosine signalling while inhibiting noradrenaline release with clonidine resulted in synapse loss (-0.8 synapses/hr) in tectal neurons (**Figure 4c**,**d**), which express both adenosine and adrenergic receptors (**Extended Data Figure 12**). These results demonstrate that daytime sleep can support synapse loss under conditions of high sleep pressure and low noradrenergic tone, possibly via direct signalling events.

**Figure 4:**
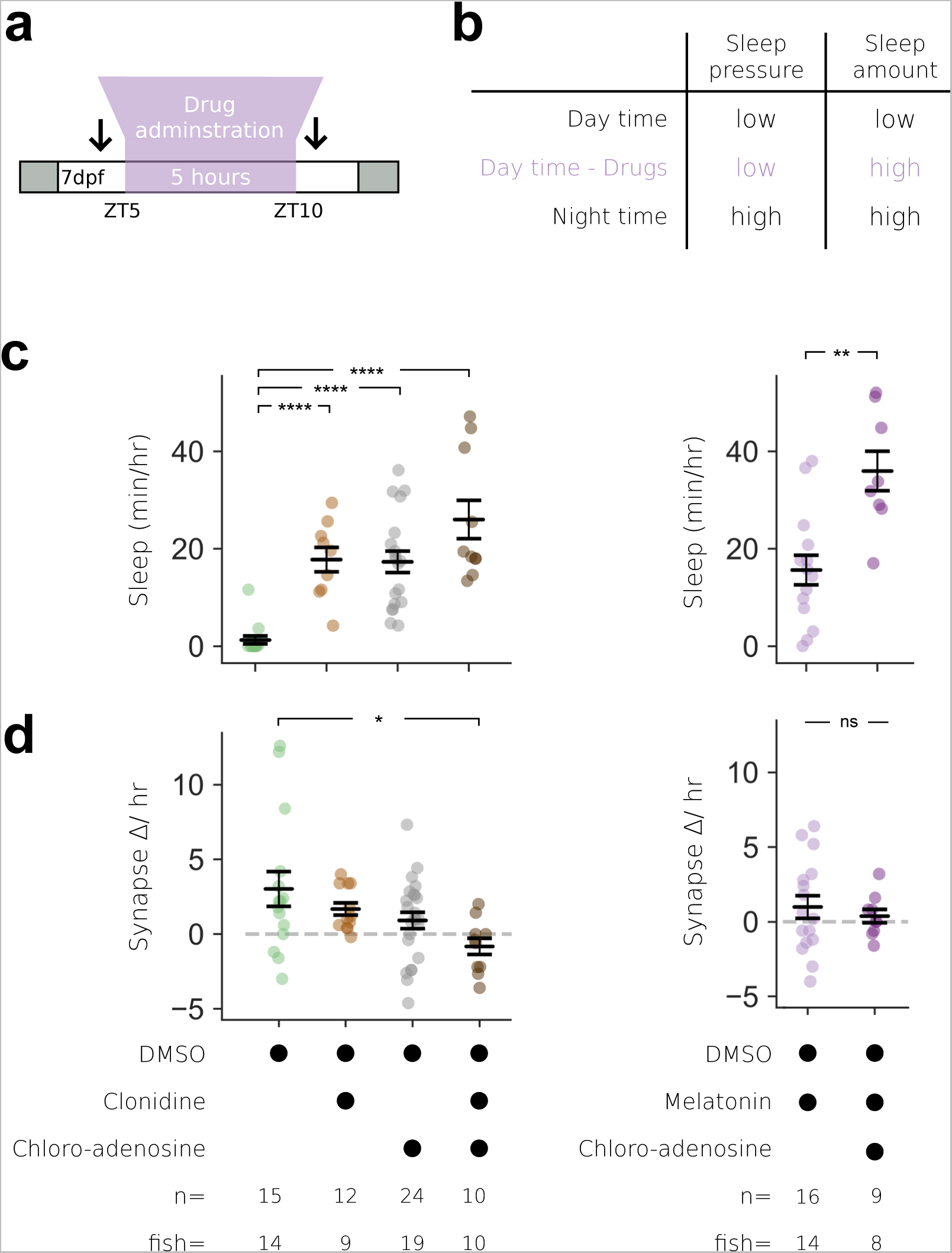
Single neuron synapse loss during sleep is driven by boosting adenosine and blocking noradrenaline. **a**, Larvae were temporarily treated with sleep promoting drugs during the day (ZT5-10). Black arrows indicate the imaging periods before and after drug treatment. **b**, Drug induced sleep during the day disentangles sleep pressure (i.e., low) from sleep amount (i.e., high), which are otherwise tightly correlated. **c**, Drug-treated larvae sleep significantly more during the day than DMSO-treated controls. **d**, During the day (from ZT5-ZT10), synapse counts increase under all control and drug conditions, except during co-administration of clonidine and 2-choloradenosine, when synapses are significantly lost. Black lines indicate mean ±SEM. *P<0.05, **P<0.01, ***P<0.001, ****P<0.0001 Kruskal-Wallis with post-hoc Dunn’s multiple comparison test.

## Discussion

The Synaptic Homeostasis Hypothesis (SHY) proposes that synapse numbers and strength increase during wake and decrease during sleep. By tracking synapses of single tectal neurons through sleep/wake states and circadian time, our data resolves several outstanding questions about the scale, universality, and mechanisms of sleep-linked plasticity. We show that SHY-concordant dynamics of the synapse population within single neurons are present on average across many cells, but when examined on a neuron-by-neuron basis more diverse patterns of synapse change are revealed. These observations may explain some discrepancies among previous studies of SHY, as these single-neuron synaptic dynamics would not be captured by population level, single time-point snapshots of synapse number or function. We also show that sleep is necessary but not sufficient for synaptic loss, since synapse loss occurred only when sleep was accompanied by high sleep pressure associated with adenosine signalling and low noradrenergic tone. Adenosine signalling has been shown to promote Homer1a-dependent downscaling and destabilisation of synapses, whereas noradrenergic signalling has been found to prevent this process^55^. Our data link these mechanisms to sleep-pressure and sleep behaviour *in vivo*. Whether single-neuron or subcellular variation in the expression or sensitivity to these synapse-regulating signals could account for the diversity of synapse dynamics remains an intriguing possibility for future work. Sleep pressure, as reflected by the density of slow wave activity in mammalian sleep, has also been linked to changes in synapses associated with learning and memory^11,56^. We find that sleep-linked synapse loss depends on molecular signals linked to high sleep pressure, and intriguingly, also mirrors slow wave activity by occurring predominantly in the early part of the sleep period^6^. This finding raises the question whether epochs of sleep associated with low sleep pressure, such as in the latter half of the night, serve additional, non-synaptic remodelling roles. If so, the evolution, persistence, and ubiquity of these different sleep epochs could be under specific regulatory and selective pressures.

## Methods

### Animals

Zebrafish husbandry and experiments were conducted following UCL Fish Facility standard protocols and under project licenses PA8D4D0E5 and PP6325955 awarded to JR, according to the UK Animal Scientific Procedures Act (1986). Embryos were kept in Petri dishes in fish water (5mM NaCl, 0.17mM KCl, 0.33mM CaCl_2_, 0.33mM MgSO_4_ and 0.1% Methylene blue) in a 14hr:10hr light:dark cycle incubator at 28°C. The sex of AB/TL zebrafish larvae is not biologically determined at the early developmental stages used for these studies.

### Cloning and transgenesis

Transgene constructs that simultaneously encode FingR targeting PSD95 and membrane markers of neuronal morphology were made using the In-Fusion HD Cloning System (Clontech). First, the GFP in a pCS2-P2A-GFP-CAAX was replaced with mKate2f by combining the linearized pCS2 (via inverse PCR) with amplified mKate2f from dUAS-mKate2f (gift from the Tada lab, UCL) with 15bp overhangs complementary to pCS2 site of insertion (**Extended Data Table 1**). Next, the template plasmid pTol2-zcUAS:PSD95.FingR-EGFP-CCR5TC-KRAB(A) (from the Bonkowsky lab, University of Utah, Addgene:72638) was linearized by inverse PCR after the KRAB(A) sequence (**Extended Data Table 1**). The P2A-mKate2f sequences were then amplified with 15bps overhangs complementary to pTol2-zcUAS:PSD95.FingR-EGFP-CCR5TC-KRAB(A) insertion site (**Extended Data Table 1**) and combined with the linearized FingR template.

To generate the stable *Tg(UAS:FingR(PSD95*)*-GFP-CCR5TC-KRAB(A*)*-P2A-mKate2f*) line, purified pTol2-zcUAS:PSD95.FingR-EGFP-CCR5TC-KRAB(A)-P2A-mKate2f DNA construct was sequenced to confirm gene insertion and co-injected (10 ng/µl) with emx3:Gal4FF^1^ (10ng/µl) and *tol2 transposase* mRNA (100 ng/µl) at 1 nl into wildtype TL embryos at the one-cell stage. At 3 dpf, injected embryos were screened for mosaic expression of the mKate2f, then raised to adulthood. The *tol2 transposase* mRNA was *in vitro* transcribed from the NotI-linearized pCS-TP6287 plasmid (gift from Wilson lab, UCL) using an SP6 mMESSAGE mMACHINE Kit (Ambion, USA). RNA was purified using RNA Clean & Concentrator Kits (Zymo Research, USA). Germline transmission was determined by mating adult fish to *nacre* mutants (*mitfa^w2^*^/*w2*^, pigmentation mutants^2^) and subsequently identifying their progeny for mKate2f fluorescence, then raising to adulthood to establish a stable *Tg(UAS:FingR(PSD95*)*-GFP-CCR5TC-KRAB(A*)*-P2A-mKate2f*)*^u541^*; *Tg(emx3:Gal4FF*)*^u542^* line. Due to the negative feedback mechanism in the system, *Tg(UAS:FingR(PSD95*)*-GFP-CCR5TC-KRAB(A*)*-P2A-mKate2f*) expression is extremely low. To increase the number of transgene copies and the level of expression in the background reporter line, the double transgenic *Tg(UAS:FingR(PSD95*)*-GFP-CCR5TC-KRAB(A*)*-P2A-mKate2f*) *; Tg(emx3:Gal4*) fish were incrossed for imaging experiments and maintained by alternating incrosses and outcrosses to *nacre* mutants.

### Whole-mount synaptic immunohistochemistry and imaging

Staining for MAGUK expression was done by whole-mount immunohistochemistry adapted from Sheets *et al* ^3^. 2 dpf zebrafish larvae were dechorionated and fixed with 4% formaldehyde methanol-free (Pierce™ Thermofisher, #28906) in BT buffer (1.0g sucrose, 18.75µl 0.2M CaCl2, topped up to 15 ml with PO_4_ buffer-- 8 parts 0.1M NaH_2_PO_4_ and 2 parts 0.1M Na_2_HPO_4_). To increase the signal-to-noise ratio, fixing time was decreased to 1.5- 2hr at 4°C, although this led to softer samples. Samples were washed with PO_4_ buffer and dH_2_O for 5 minutes at room temperature (RT), then permeabilized with ice-cold 100% acetone for 5 minutes at -20°C. After washing with dH_2_O and PO_4_ buffer for 5 minutes each, specimens were blocked with blocking buffer containing 2% goat serum, 1% bovine serum albumin (BSA) and 1% dimethyl sulfoxide (DMSO) in 0.1 M Phosphate buffered saline (PBS) pH 7.4 for at least 2 hours. The samples were then incubated with primary antibodies (see below for list) diluted in blocking buffer at 4°C overnight. Embryos were washed 4-6 times for at least 20 minutes in blocking buffer at RT and incubated in secondary antibodies overnight at 4°C. To remove unbound secondary antibodies, the embryos were washed again and transferred to glycerol in a stepwise manner up to 80% glycerol in PBS.

The primary antibodies used for staining were Anti-pan-MAGUK (mouse monoclonal, clone K28/86, Millipore) and Anti-tRFP (rabbit polyclonal, AB233, Evrogen), both at 1:500 dilution. To avoid over-amplification of signal outside of the synapse, FingR(PSD95)-GFP puncta were visualized using its own fluorescence. The following secondary antibodies were used at 1:200 dilution; Alexa-Fluor 568 goat anti-rabbit IgG; and Alexa-Fluor 633 goat anti-mouse IgG monoclonal (Life Technologies).

Confocal images were obtained using a Leica TCS SP8 system with HC PL APO 20x/0.75 IMM CS2 multi-immersion objective set to glycerol (Leica Systems). Z-stacks were obtained at 1.0μm depth intervals with sequential acquisition settings of 1024 x 1024 pixels. The raw images were compiled using NIH Image J software (http://imagej.nih.gov/ij/). To analyse the colocalization of the puncta, maximum projections of 5-10μm were taken for each cell. Grey values were taken from the cross-section of the puncta using the *plot-profile* tool from ImageJ. Puncta grey values were normalized against the whole stack grey value of their respective channels.

The colocalization and relationships between FingR(PSD95)-GFP and antibody staining were analysed using custom written scripts on Python (available at https://github.com/anyasupp/single-neuron-synapse). For colocalization of FingR and antibody puncta (and vice versa), the presence of puncta with maximum normalized grey value of at least 50% higher than the baseline) were used. To estimate the size of the puncta, the normalized grey values were fitted with a non-gaussian prior to finding the full width half maximum (FWHM).

### Single-cell FingR(PSD95) expression using electroporation

To sparsely label single tectal cells a FoxP2.A:Gal4FF activator plasmid (gift from Martin Meyer, King’s College London) was electroporated into the *Tg(UAS:FingR(PSD95*)*-GFP-ZFC(CCR5TC*)*-KRAB(A*)*-P2A-mKate2f*)-positive larvae at 3 dpf following the method of Nikolaou and Meyer (2015)^4^. Anaesthetized 3 dpf zebrafish larvae were mounted in 1% low-melting point agarose (Sigma), perpendicular to a glass slide in a Petri dish filled with electroporation buffer (180mM NaCl, 5mM KCl, 1.8mM CaCl2, 5mM HEPES, pH 7.2) with 0.02% Tricaine (MS-222, Sigma-Aldrich). Excess agarose along the larval body was then removed to allow access for the electroporation electrodes. A FoxP2.A:Gal4FF construct (500 ng/μl) was injected into the midbrain ventricle together with *tol2* mRNA (20ng/μl) and Phenol-red (∼0.025%) at 5-8nL using a micro glass needle (0.58mm inside diameter, Sutter Instrument, Germany, BF100-58-15) pulled using a micropipette puller (Model P-87 Sutter Instrument, Germany). Following injection, the positive electroporation electrode was placed lateral and slightly dorsal to the hemisphere of the target optic tectum, and the negative electrode was placed lateral and ventral to the contralateral eye. Five 5ms trains of 85 V voltage pulses at 200 Hz were delivered through the electrodes using an SD9 stimulator (Grass Instruments). Electroporated larvae were screened for sparse, single-cell expression of FoxP2:FingR(PSD95)+ neurons using a 20x water-immersion objective and an LSM 980 confocal microscope with Airyscan 2 (Zeiss) at 5-6 dpf.

### Repeated Imaging of FingR-labelled synapses

For synapse tracking experiments, *Tg(UAS:FingR(PSD95*)*-GFP-CCR5TC-KRAB(A*)*-P2A-mKate2f*) larvae that were electroporated with FoxP2.A:Gal4FF were reared at 28°C under various light schedules (**Extended Data Table 2**). At 5-6 dpf, larvae were visually screened for the expression of single or sparsely labelled FoxP2.A:FingR(PSD95)+ neurons in the tectum using a 20x water-immersion objective and an LSM 980 confocal microscope with Airyscan 2 (Zeiss) and placed into individual wells of a 6-well plates (Thermo Fisher Scientific) to keep track of individual larvae and the corresponding labelled neurons, each well containing approximately 10mL of fish water. For repeated live imaging of reticulospinal neurons, *Tg(UAS:FingR(PSD95*)*-GFP-CCR5TC-KRAB(A*)*-P2A-mKate2f*) were crossed to a Tg(*KalTA4^u508^*) driver line^5^ (gift from the Bianco lab at UCL) and visually screened for larvae that had the reticulospinal population labelled. For imaging FingR(PSD95)-GFP puncta, the larvae were anaesthetized with 0.02% Tricaine for 5-10 minutes and immobilized in 1.5-2% low-melting point agarose (Sigma) in fish water. The larvae were head-immobilized with the tail free and allowed to recover from anaesthesia during imaging. Imaging was performed at the appropriate Zeitgeber/circadian time (ZT, where ZT0=lights ON) according to the experimental paradigm (**Extended Data Table 2**). For day:night synapse tracking, larvae were repeatedly imaged approximately at ZT0-2 and ZT10-12 at 7 dpf, 8 dpf, and 9 dpf at 28.5°C with chamber lights ON. Imaging performed during the dark phase (ZT14-24) were kept at 28.5°C with the chamber lights OFF. When immobilizing the larvae for night imaging, the handling was performed under dim red light (Blackburn Local Bike Rear Light 15 Lumen, UK). After imaging, larvae were unmounted from agarose by releasing agarose around their heads and allowing larvae to independently swim out of the agarose. Unmounted larvae were then placed back into individual wells of 6- well plates.

FingR(PSD95)+ neuron image stacks were acquired using a 20x water-immersion objective and an LSM 980 confocal microscope with Airyscan 2 (Zeiss). GFP and mKate2f were excited at 488nm and 594nm, respectively. Z stacks were obtained at 0.34μm voxel depth with sequential acquisition settings of 2024 x 2024 pixels (0.0595376 x 0.0595376μm pixel width x height) and 16-bit using SR4 mode (imaging 4 pixels simultaneously). Pixel alignment and processing of the raw AiryScan stack were performed using ZEN Blue software (Zeiss).

### Locomotor activity assay

Tracking of larval zebrafish behaviour was performed as previously described^6^, with slight modifications. Zebrafish larvae were raised at 28.5°C on 14hr:10hr light:dark cycle or according to the needs of the experimental design (**Extended Data Table 2**). At 5-6 dpf each FoxP2.A:FingR(PSD95)+ larva was placed into individual wells of a 6-well plate (Thermo Fisher Scientific) containing approximately 10mL of fish water. Locomotor activity of some larvae was monitored using an automated video tracking system (Zebrabox, Viewpoint LifeSciences) in a temperature-regulated room (26.5°C) and illuminated with white lights on either 14hr:10hr L:D cycles or constant light conditions at 480-550 lux with constant infrared illumination. The larval movement was recorded using the Videotrack ‘quantization’ mode with the following detection parameters: detection threshold, 15; burst, 100; freeze, 3; bin size, 60s. The locomotor assay data were analyzed using custom MATLAB (MathWorks) scripts available at https://github.com/JRihel/Sleep-Analysis. Any one-minute period of inactivity was defined as one minute of sleep, according to established convention for larval zebrafish^7^. Experiments examining the effects of drug treatment on behaviour that did not involve live imaging, such as the clonidine dark pulse experiment (**Extended Data** Figure 11), 24-well (Thermo Fisher Scientific) and 96-well plates (Whatman) were used instead of the 6-well plates used for synapse imaging experiments. Sleep latency for **Extended Data** Figure 9a-c was calculated using frame-by-frame data (collected at 25 fps), using code available at (https://github.com/francoiskroll/FramebyFrame).

### Sleep deprivation assay

Zebrafish larvae were raised at 28.5°C on 14hr:10hr light:dark cycle to 6 dpf, when they were videotracked (see Locomotor activity assay). Randomly selected 7 dpf larvae were then sleep deprived for 4 hours immediately after lights off from ZT14-18. Non-deprived control larvae were left undisturbed. Larvae that were individually housed in 6-well plates were manually sleep deprived under dim red light (Blackburn Local Bike Rear Light 15 Lumen, UK) by repeated gentle stimulation using a No. 1-2 paintbrush (Daler-Rowney Graduate Brush, UK) to prevent larvae from being immobile for longer than 1 minute. For most stimulations, this required only putting the paintbrush into the water; if larvae remain immobile, they were gently touched. The 4hr sleep deprivation protocol was performed by experimenters in 2 hr shifts. All sleep deprived and control larvae were imaged at ∼ZT14 and ZT18 on 7 dpf and again at ZT0 on 8 dpf (see **Repeated imaging of FingR-labelled synapses**).

### Drug exposure for live imaging

*Tg(UAS:FingR(PSD95*)*-GFP-CCR5TC-KRAB(A*)*-P2A-mKate2f*) larvae that had been electroporated with FoxP2.A:Gal4FF (see **Single-cell FingR(PSD95) expression using electroporation**) were kept on a 14hr:10hr L:D cycle until 7 dpf, then imaged at ZT4-5 (see **Repeated imaging of FingR-labelled synapses**). Larvae were transferred to individual wells of a 6-well plate containing 10mL of sleep promoting drugs, alone or in combination, as follows: 30µM melatonin (M5250, Sigma) in 0.02% DMSO; 30µM of clonidine hydrochloride (C7897, Sigma) in 0.02% DMSO; 45µM 2-Chloroadenosine (C5134, Sigma) in 0.02% DMSO; or 0.02% DMSO in fish water as controls^6,8–10^. Combinations of drugs were applied at the same concentrations as the single dose conditions, maintaining the final DMSO concentration of 0.02%. Sleep induction was monitored with videotracking (see **Locomotor activity assay**) for 5 hrs, after which drugs were removed by 2-3 careful replacements of the fish water using a transfer pipet followed by transferring the larvae individually to a new 6- well plate with fresh water. Larvae were then re-imaged using AiryScan (see **Repeated imaging of FingR-labelled synapses**).

### Tectal cell segmentation and clustering

The morphology of tectal neurons at 7 dpf was segmented and measured using Imaris 8.0.2 software (Bitplane) and ImageJ (NIH). The total filament length for each neuron was obtained using the Imaris *Filament* function. The anterior-posterior (AP) span of the distal arbour was calculated using the *Measurement* function at an orthogonal view in 3D. The relative proximal arbour locations were calculated by dividing the proximal arbour distance from the nucleus by the total length of the neuron obtained using *Filament* function on Imaris. The distance from the skin, distal arbour thickness, and distal arbour to skin distance were obtained using the rectangle *Plot_Profile* tool on ImageJ at an orthogonal view of the neuron to calculate the fluorescence intensity across the tectal depth. The intensity profiles were then analysed using custom Python scripts to obtain the maximum width using area under the curve functions following the published methods of ^4,11^.

Additional clustering and statistical analyses were performed using custom written scripts written in Python (available at https://github.com/anyasupp/single-neuron-synapse). For segmentation clustering, six morphological features of FoxP2.A cells were standardized and reduced in dimensionality by projecting into principal component analysis (PCA) space. The first four components, which explained 89% of the variance, were selected to use for clustering. These components were then clustered using K-means with K ranging from 1 to 11. Using the elbow method, Calinski Harabasz coefficient, and silhouette coefficient found k = 4 to be the optimal number of k clusters.

### Puncta quantification and statistics

All image files of synapse tracking experiments were blinded by an independent researcher prior to segmentation and puncta quantifications. To count number of FingR(PSD95)-GFP puncta, each neuron’s morphology was first segmented using the *Filament* function in Imaris 8.0.2 software (Bitplane). FingR(PSD95)-GFP puncta were labelled using the *Spots* function, thresholded using the *Quality classification* at approximately 130-200 depending on the image file. The number and location of GFP puncta were also manually checked for accuracy. FingR(PSD95)-GFP puncta lying on the FingR+ neuron (mKate2f red channel) were extracted using the *Find Spots Close to Filamen*t XTension (add-on in IMARIS). The average 3D nuclear intensity per neuron per time point was obtained using *Spots* function on Imaris.

The percentage change in synapse number and intensity were calculated by the following formula:

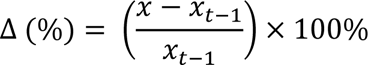

Where *x* represents either synapse number or intensity and *x*_*t*−1_ is the respective synapse number or intensity at the previous time point. Mixed-designed ANOVA (mixed-measure ANOVA), post-hoc pairwise t-tests, and Student’s t-test were implemented using Python^12^. Values in figures represent the average ±68% CI unless stated otherwise.

Synapse intensity was calculated using the ratio of the normalized average FingR(PSD95)- GFP intensity and mKate2f, to account for depth-dependent signal reduction^13^. First, the average FingR(PSD95)-GFP and mKate2f (cell morphology) intensities at the same location within the neuron were extracted using the Imaris *Spots* function. Next, these average intensity values were normalized with their respective channel maximum and minimum value to account for larval position inconsistencies between imaging using:

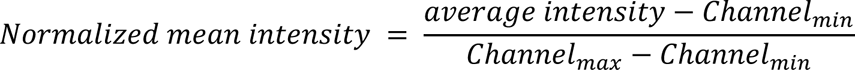

Depth-dependent signal reduction was corrected by calculating the FingR(PSD95)- GFP:mKate2f ratio using:

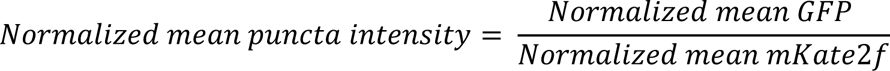

### *per3* circadian rhythm bioluminescence assay

6 dpf larvae from a *Tg(per3:luc*)*^g1^; Tg(elavl3:EGFP*)*^knu3^* incross were individually placed in wells of 24-well plates in water containing 0.5 mM beetle luciferin (Promega). From ZT14 (the light to dark transition) the following day, half of the larvae were subjected to a sleep deprivation paradigm (see **Sleep deprivation assay**) under dim red light, while the others were left undisturbed in similar lighting conditions. At the end of the 4-hour sleep deprivation period, the larvae were individually transferred to the wells of a white-walled 96-round well plate (Greiner Bio-One) and sealed with an oxygen-permeable plate-seal (Applied Biosystems). Bioluminescence photon counts, reflecting luciferase expression driven by the *per3* promoter, were sampled every 10 minutes for three consecutive days, in constant dark at 28°C, using a TopCount NXT scintillation counter (Packard).

### Hybridization Chain Reaction (HCR) fluorescence in situ hybridization

FoxP2.A neurons were sparsely labelled with GFP by co-electroporating wildtype AB larvae with FoxP2.A:Gal4FF and UAS:EGFP^1^ at 500ng/µl each (see **Single-cell FingR(PSD95) expression using electroporation**). Whole-mount HCR was performed on larvae with FoxP2.A neurons positive for GFP at 7 dpf using an adapted protocol from Choi et al (2018)^14^. Briefly, larvae were fixed with 4% PFA, 4% sucrose overnight at 4°C. The following day larvae were washed with PBS to stop fixation and brains were removed by dissection. The dissected specimens were permeabilized using proteinase K (30µg/ml) for 20 minutes at RT, then washed 2x in PBS with 0.1% Tween (PBST), before being post-fixed in 4% PFA for 20 minutes at RT. Larvae were then washed in 0.1% PBST and prehybridized with prewarmed HCR hybridization buffer (Molecular Instruments, USA) for 30 minutes at 37°C.

Probes targeting multiple genes associated with different types of adenosine or adrenergic receptors were combined and labelled to the same hairpins. For example, probes detecting *adora1a-b* (encoding for adenosine receptor A1a and A1b) contains initiators that correspond with hairpins (B3) labelled with Alexa 546 fluorophore, whereas *adora2aa*,*-ab*,*-b* (encoding for adenosine receptors A2aa, A2ab, and A2b) contains initiators that correspond with hairpins (B5) labelled with Alexa 647 fluorophore (see **Supplementary File 1**). Probe solutions consisting of cocktails of HCR probes for each transcript (Thermo Fisher Scientific, UK) were prepared with a final concentration of 24nM per HCR probe in HCR hybridization buffer. Larvae were then incubated in probe solutions overnight at 37°C. Excess probes were removed by washing larvae 4 x 15 minutes with probe wash buffer (Molecular Instruments, USA) at 37°C followed by 2 x 5 minutes of 5x SSCT buffer (5x sodium chloride sodium citrate and 0.1% Tween) at RT. Preamplification was performed by incubating samples with amplification buffer (Molecular Instruments, USA) for 30 minutes at RT. Hairpin h1 and hairpin h2 were prepared separately by snap cooling 4µl of 3µM stock at 95°C for 20 minutes and 20°C for 20 minutes. Larvae were then incubated with h1 and h2 hairpins in 200µL amplification buffer overnight in the dark at RT. Excess hairpins were washed thoroughly the next day with 2 x 5 minutes and 3x 30 mins of SSCT at RT. Specimens were then imaged using a 20x water-immersion objective and an LSM 980 confocal microscope with Airyscan 2 (Zeiss). Endogenous GFP signal from FoxP2.A were visualized without amplification.

### Data availability

Data and code can be found https://github.com/anyasupp/single-neuron-synapse. Sleep analysis code https://github.com/JRihel/Sleep-Analysis. Frame by frame analysis code can be found at https://github.com/francoiskroll/FramebyFrame.

## Acknowledgements

We thank all current and past members of the Rihel lab for helpful discussions and feedback on this project and the zebrafish community for sharing protocols and reagents; special thanks to Sumi Lim (UCL) for her assistance in blinding experimental files, Alexandra Gilbert (UCL) for help with early sleep deprivation experiments, Lavinia Sheets (Washington University) for her guidance on synapse immunohistochemistry, Leah Elias (Johns Hopkins University) for her guidance on gene expression, Nikolas Nikolaou (University of Bath) for sharing his knowledge of FoxP2.A neurons, Chintan Trivedi (UCL) for his help with designing HCR probes, UCL Fish Facility for fish husbandry and UCL Imaging Facility for their expertise. This work was supported by UCL Research Scholarship (to AS), an EMBO Fellowship awarded to DGL (ALTF 1097-2016), a Medical Research Council studentship (MR/W006774/1 to EB), a European Research Council Starting Grant (282027 to JR), and a Wellcome Trust Investigator Award (217150/Z/19/Z to JR)

## Author Contributions

JR and AS conceived of and designed all experiments with input from DGL. AS performed all experiments with help from DGL (circadian clock/dark pulse), EB (HCR), DGL and JR (sleep deprivation). AS, DGL, and JR wrote the manuscript with input from EB.

## Additional Information

Supplementary Information is available for this paper. Correspondence and requests for materials should be addressed to Jason Rihel.

## Extended Data Figures

**Extended Data Figure 1.**
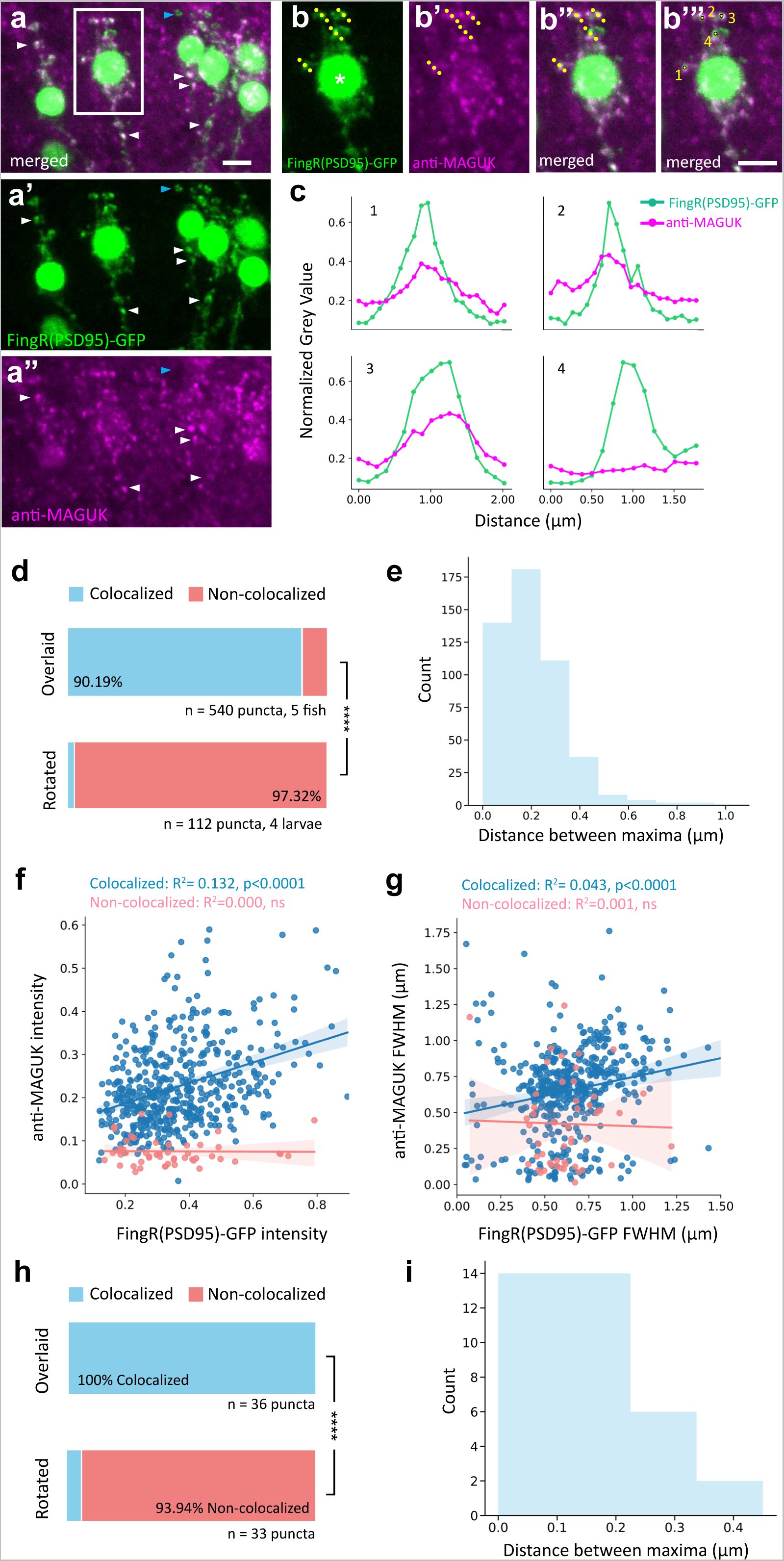
The modified FingR(PSD95)-GFP construct labels synapses *in vivo*. **a-a ’’**, Maximum projection (Z-stack, ∼10 μm) of anti-MAGUK immunohistochemistry and endogenous fluorescence of FingR(PSD95)-GFP in the spinal cord of 2 dpf *Tg(mnx1:Gal4*) larvae. Examples of FingR(PSD95)+ puncta co-labelled by anti-MAGUK are indicated by white arrowheads; an example of a FingR(PSD95)+ not labelled by anti-MAGUK is indicated by the blue arrowhead. Scale bar, 5μm. **b-b’’’**, Higher magnification (white box from **a**) depicting how sectional grey values for each synapse were obtained. **b**, The FingR(PSD95)-GFP channel showing part of a neuron with its nucleus (asterisk) and synaptic puncta (green). Dotted lines indicate example cross-sectional areas obtained for each synapse. **b**’, Anti-MAGUK puncta of the same neuron. **b’’**,**b’’’**, FingR(PSD95)-GFP and MAGUK channels merged, with examples of cross-sections 1-4 that are depicted in **c**. Scale bar, 5μm. **c**, Examples of normalized cross-sectional grey values for anti-MAGUK signals and FingR(PSD95)-GFP signal for the same puncta (numbered 1-4 in **b’’’**). Three examples in which FingR(PSD-95)-GFP co-localized with anti-MAGUK signals (#1-3) and one example (#4) where a FingR(PSD-95)-GFP punctum did not co-localize with MAGUK. See **Methods** for details. **d**, Percentage of FingR(PSD-95)-GFP synapses that co-localized with anti-MAGUK+ puncta (blue). As a control for chance co-localization, the calculation was repeated on images in which the anti-MAGUK image was rotated by 90° relative to the FingR(PSD-95)-GFP channel. ****P< 0.0001 Chi-square. **e**, Histogram of the distance between all co-localized FingR(PSD95)-GFP and anti-MAGUK cross-sectional grey value peaks. **f-g**, The intensity and Full Width Half Max (FWHM) of FingR(PSD95)-GFP and anti-MAGUK puncta are weakly, but significantly, positively correlated. Blue and red lines depict the linear regression curve and 95% CI for the colocalized and non-colocalized populations, respectively. n= 540 puncta, 5 fish (data as in **d**). **h**, Percentage of anti-MAGUK+ puncta that co-localized with FingR(PSD-95)-GFP synapses (blue). As a control for chance co-localization, the calculation was repeated on images in which the FingR(PSD-95)-GFP image was rotated by 90° relative to the anti-MAGUK channel. ****P<0.0001 Chi-square. **i**, Histogram of the distance between co-localized anti-MAGUK and FingR(PSD95)-GFP cross-sectional grey value peaks.

**Extended Data Figure 2:**
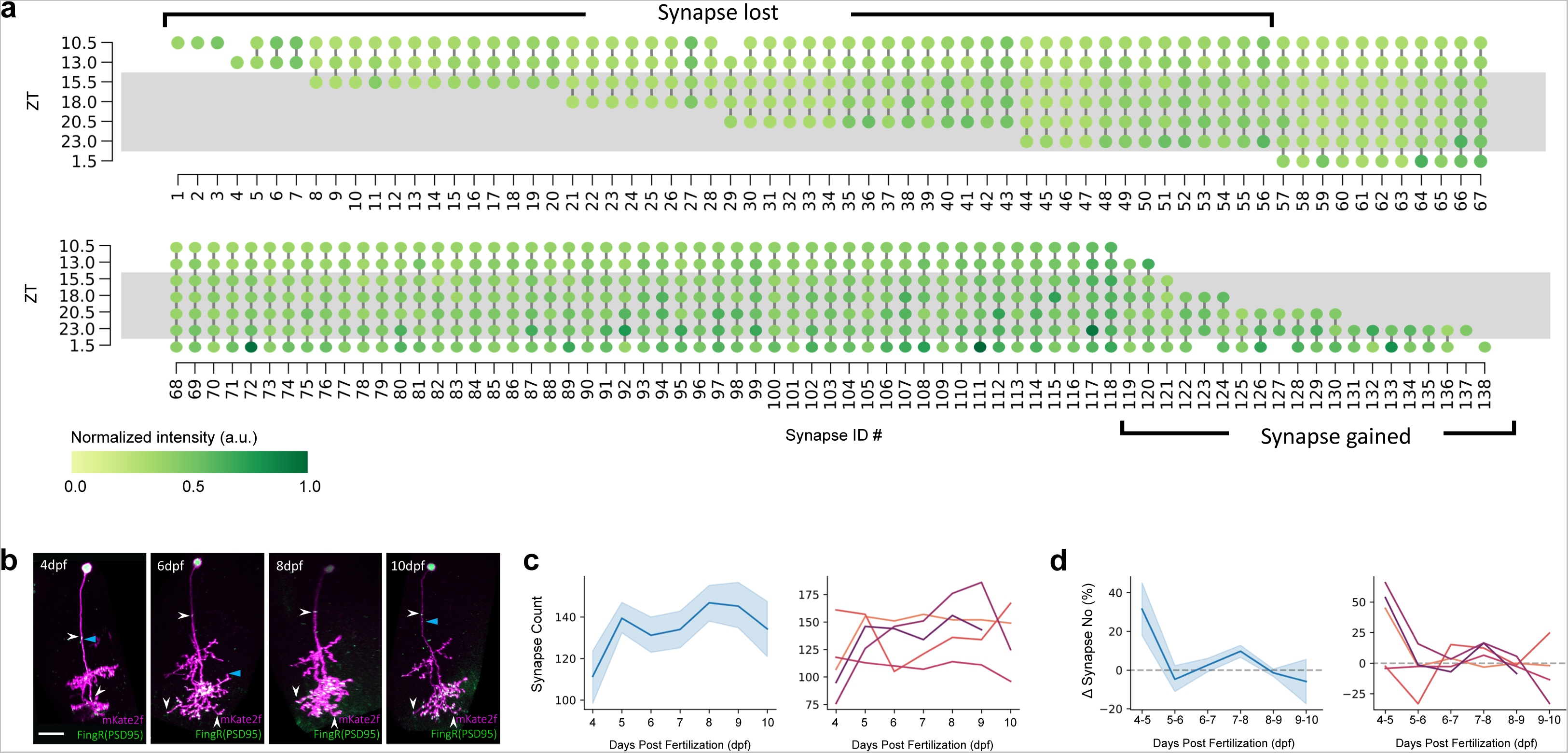
The synapse number of single tectal neurons is developmentally stable at 6-9 dpf. **a**, The full map of synapse tracking from the neuron in Figure 1c. Each column depicts a synapse, and the colour indicates the normalized GFP intensity of each synapse. In this example, 56 synapses disappeared and 20 synapses appeared during the imaging, resulting in a net change of -36 synapses. Grey bars depict night (ZT14-24). **b**, Example of a single FoxP2.A:FingR(PSD95)+ neuron imaged through development from 4-10 dpf. Nuclei and synapses are FingR(PSD95)-GFP+ (green), and cellular morphology is labelled by mKate2f (magenta). White arrowheads indicate examples of puncta that persisted through time. Blue arrowheads indicate examples of synapses gained/lost through time. Scale bar, 15μm. **c**, Synapse counts across all neurons (average and 68% CI) (**left**) and for single neurons through 4-10 dpf (**right**). **d**, The average percentage change in synapse number and 68% CI calculated from the previous time point (**left**) and for each neuron (**right**). The percentage change in synapse number across time is close to zero between 6-9 dpf. n= 5 cells, 5 larvae.

**Extended Data Figure 3:**
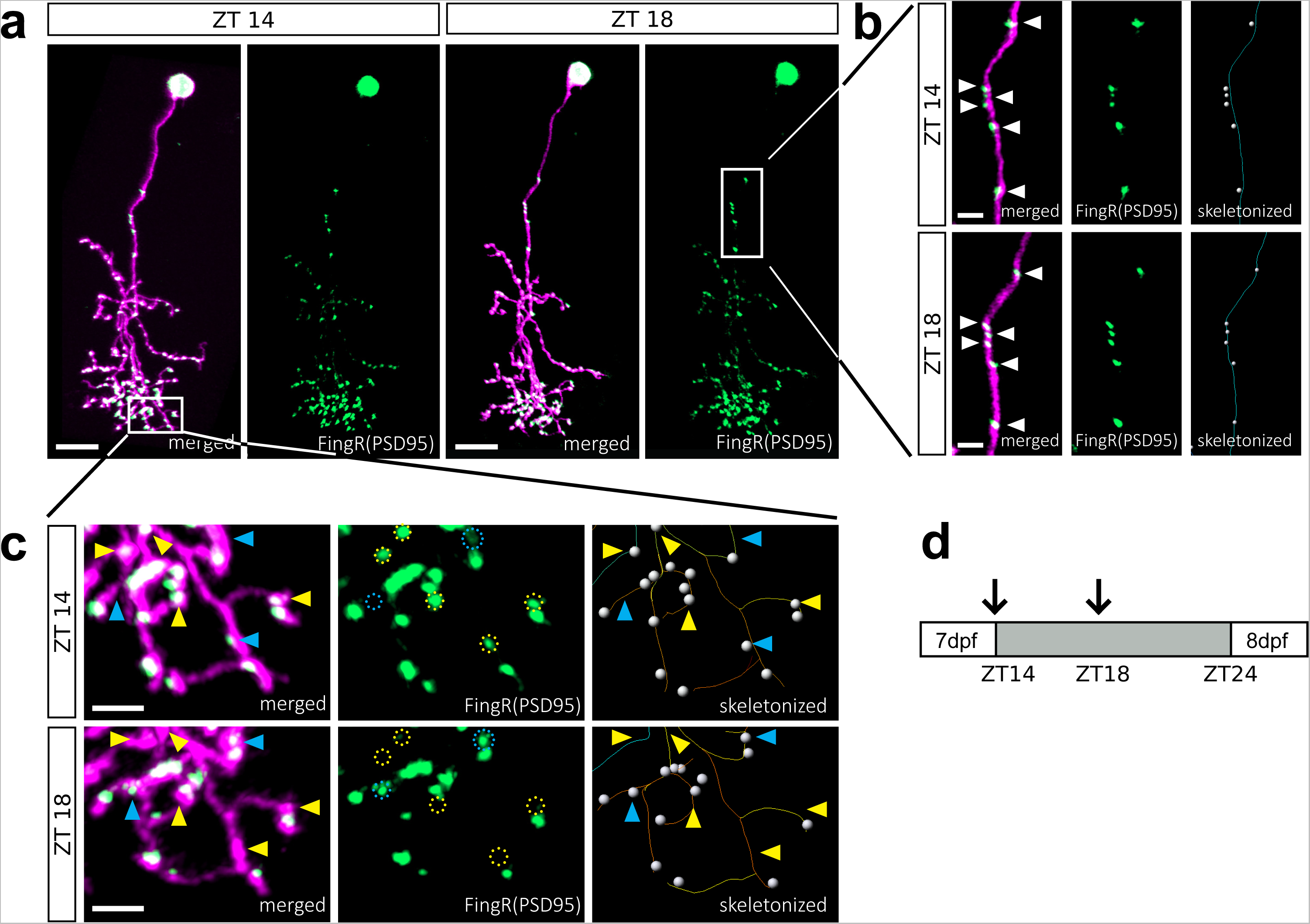
Example of a single FoxP2.A:FingR(PSD95)+ neuron at ZT14 and ZT18. **a**, A single FoxP2.A:FingR(PSD95)+ tectal neuron imaged at ZT14 and ZT18. Nuclei and synapses are FingR(PSD95)-GFP+ (green), and cellular morphology is labelled by mKate2f (magenta). Scale bar, 10μm. **b**, Higher magnification of the primary dendrite segment (white box in **a**). Right panels show semi-automatic skeletonization (lines) of neurites and detection of FingR(PSD95)-GFP puncta (grey spheres, **Methods**). **c**, Higher magnification of a section of the distal arbour (white box in **a**). FingR(PSD95)-GFP+ puncta that appeared (blue circles and arrowheads) and disappeared (yellow circles and arrowheads) between ZT14 and ZT18 can be observed. Scale bars of **b**,**c**, 2.5μm. **d**, Schematic showing imaging times (black arrows) at ZT14 and ZT18 on the night of 7 dpf.

**Extended Data Figure 4:**
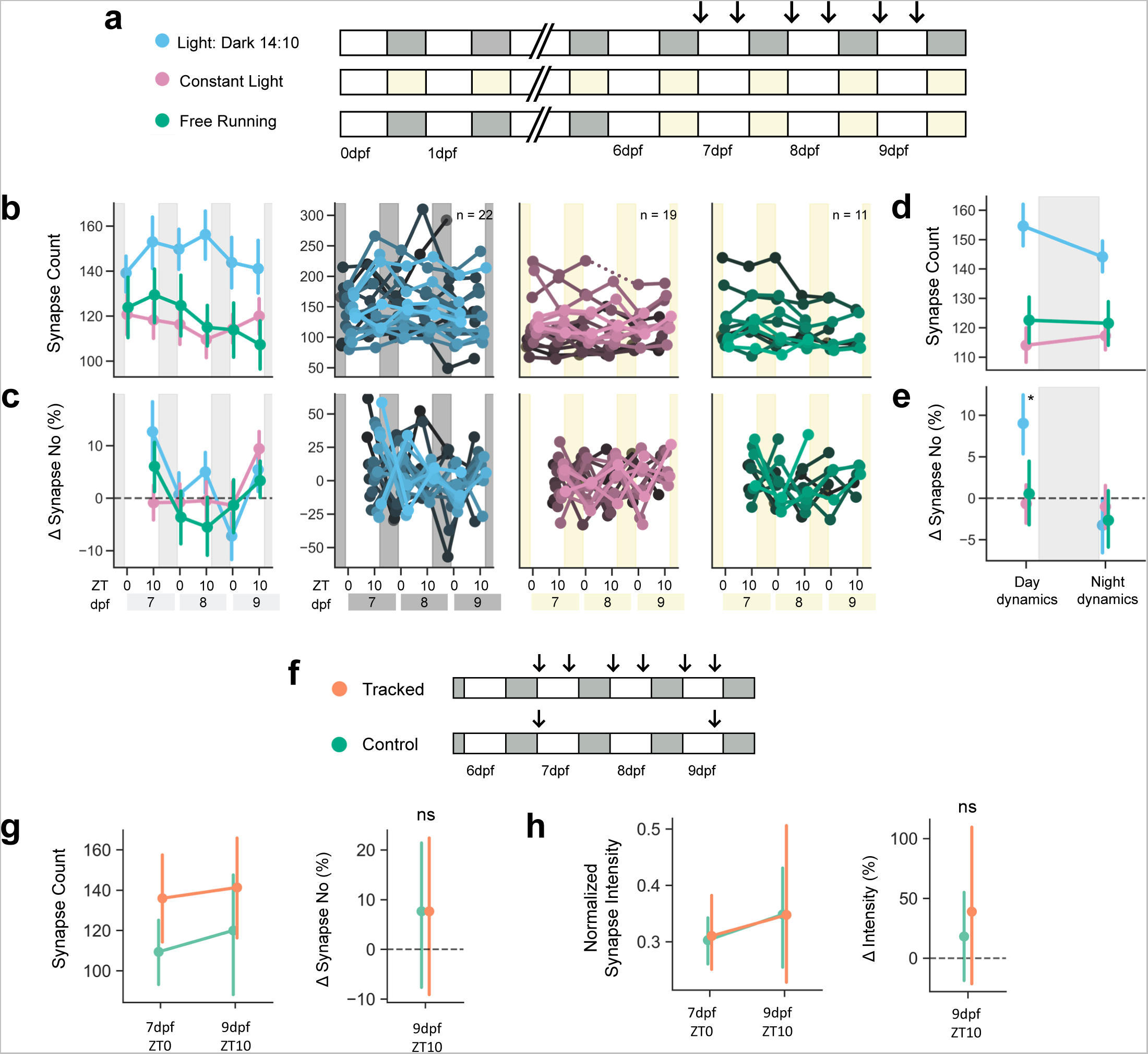
Extended tracking of single neurons over multiple days. **a**, Larvae were raised on 14hr:10hr LD cycles (blue), on constant light (pink), or switched from LD to LL at 6 dpf (‘free running’, FR, green). The arrows show the times of synapse imaging at ZT0 and ZT10 for each day from 7-9 dpf. **b**, The average and 68% CI for synapse counts at each timepoint in LD (blue), LL (pink), or FR (green) conditions from 7-9 dpf (**left**). Synapse counts for each neuron are plotted as a single line (**right**). **c**, The percentage change (average and 68% CI, **left**; each neuron, **right**) of synapse counts calculated within each neuron across time (from **b**). **d-e**, The average synapse counts and percentage change for ZT0 and ZT10 combined across all tracked days for each lighting condition (LD, LL, and FR). The ZT10 timepoint from 9 dpf was excluded to avoid interference from a new developmental round of synaptogenesis. Larvae raised in LD had a significantly higher average percentage change during the day phase than larvae raised in LL during the day phase. *P<0.05; mixed ANOVA with pairwise t-test. **f**, Schematic of experiment set up to test whether repeated imaging affected total synapse number and strength measurements. Larvae were raised in LD (indicated by white and grey boxes) and either imaged six times between 7-9 dpf at ZT0 and ZT10 each day (Tracked, orange) or imaged at the first time point ZT0 on 7 dpf and the last time point ZT10 on 9 dpf (Control, green). **g**, Average synapse counts and 68% CI at the first and last time point (7 dpf ZT0 and 9 dpf ZT10) for tracked and control larvae (**left**). The percentage changes in synapse number were not statistically different between tracked and control larvae (**right**). **h**, The normalized average synapse intensity for tracked and control larvae at the first and last time points (**left**). The percentage change in normalized average synapse intensity was not statistically different between tracked and control larvae (**right**). Controls: n=6 neurons, 4 larvae; Tracked: n=14 neurons, 14 larvae. ns, P>0.05 Student’s t-test.

**Extended Data Figure 5:**
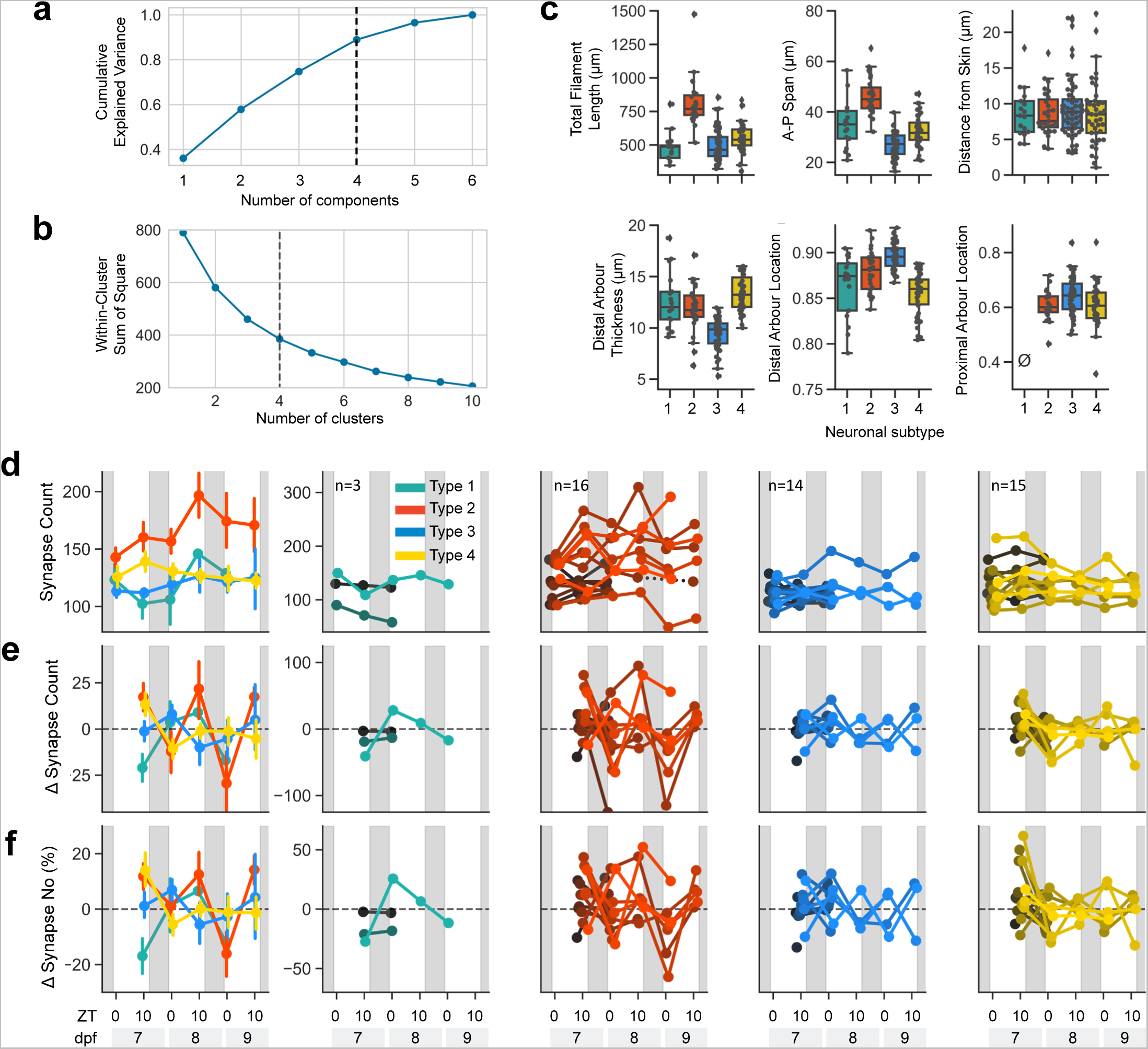
FoxP2.A tectal neurons have four morphological subtypes. **a**, Principal component analysis using the subtype morphological features depicted in Figure 2a. Four principal components (dotted line) account for >85% of the variance. **b**, The optimal number of clusters for k-means clustering was determined using the elbow method by plotting the within-cluster sum of squares. Four clusters were chosen (dotted line). **c**, The six features used to cluster FoxP2.A neurons by morphological subtype. Boxes depict the median and interquartile range and the whiskers represent the distribution for each parameter. The slashed zero means the feature is absent. **d-f**, Synapse dynamics in different FoxP2.A tectal neuron subtypes of larvae raised in normal LD conditions. **d**, Average synapse number and 68% CI of each subtype (**left**) and the puncta count for each neuron, grouped by subtype (**right panels**). **e**, Average change in synapse numbers per neuron within each subtype (**left**) and for each individual neuron, grouped by subtype (**right panels**). **f**, Average percentage change of synapse number for each subtype (**left**) and for each neuron, grouped by subtype (**right panels**).

**Extended Data Figure 6:**
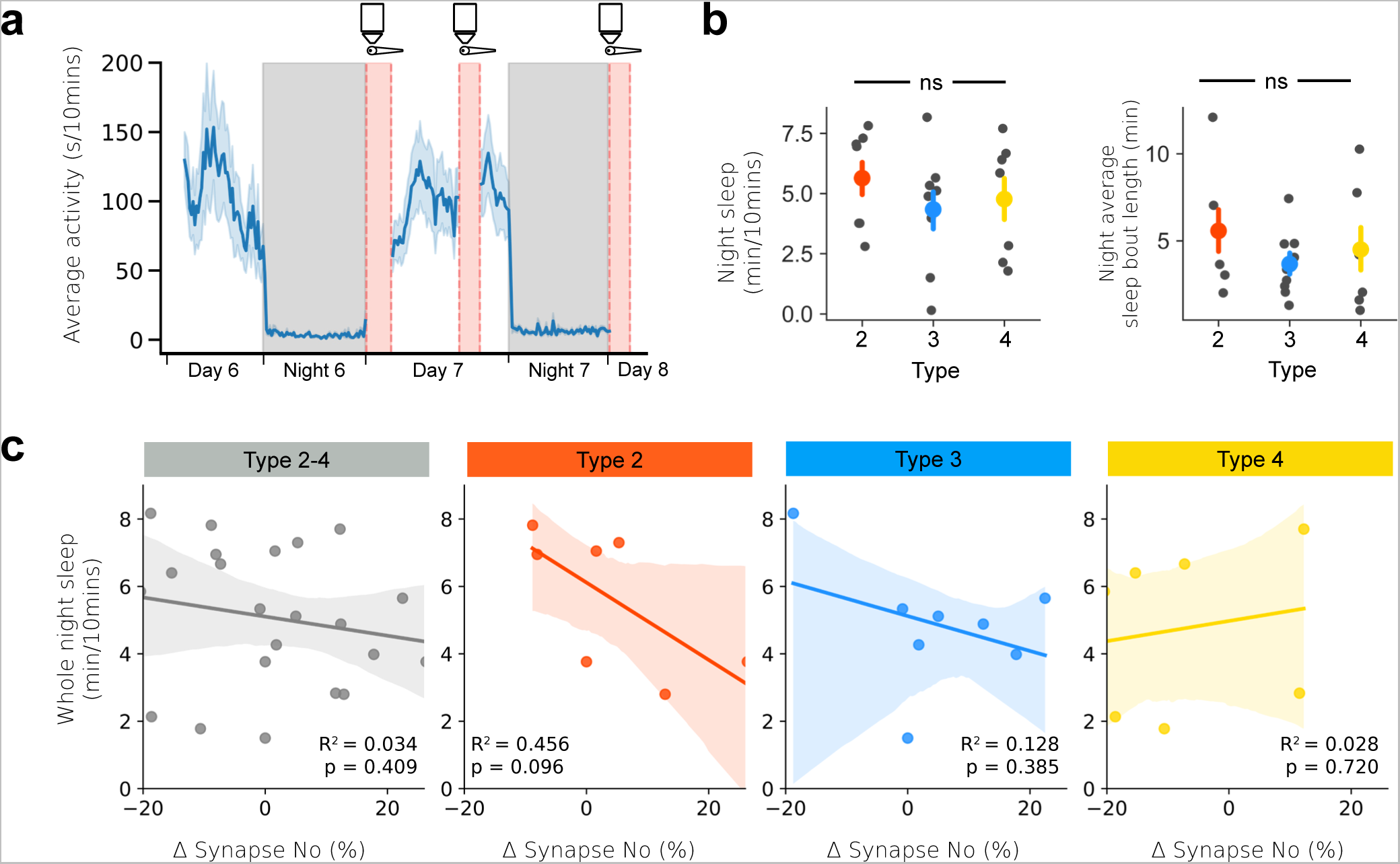
There was no systematic bias in sleep amount for larvae labelled with specific FoxP2.A tectal neurons subtypes. **a**, Schematic of behavioural and synapse tracking experiment set up. Larval locomotor behaviour was tracked on a 14hr:10hr LD cycle from 6-8 dpf. The average activity (±68% CI) of 10 example larvae are plotted across two days and nights. Larvae were removed from the tracking arena and imaged at lights on (ZT0) and again at ZT10 (dotted red bars). White and grey boxes indicate day and night periods, respectively. **b**, 7 dpf Larvae had similar levels of sleep and sleep bout lengths at night regardless of the FoxP2.A tectal neurons subtype labelled in each larva. **c**, For each neuron/larva, the average percentage change of synapse number is plotted versus the average 7 dpf night-time sleep. Linear regression is fitted with 95% CI.

**Extended Data Figure 7:**
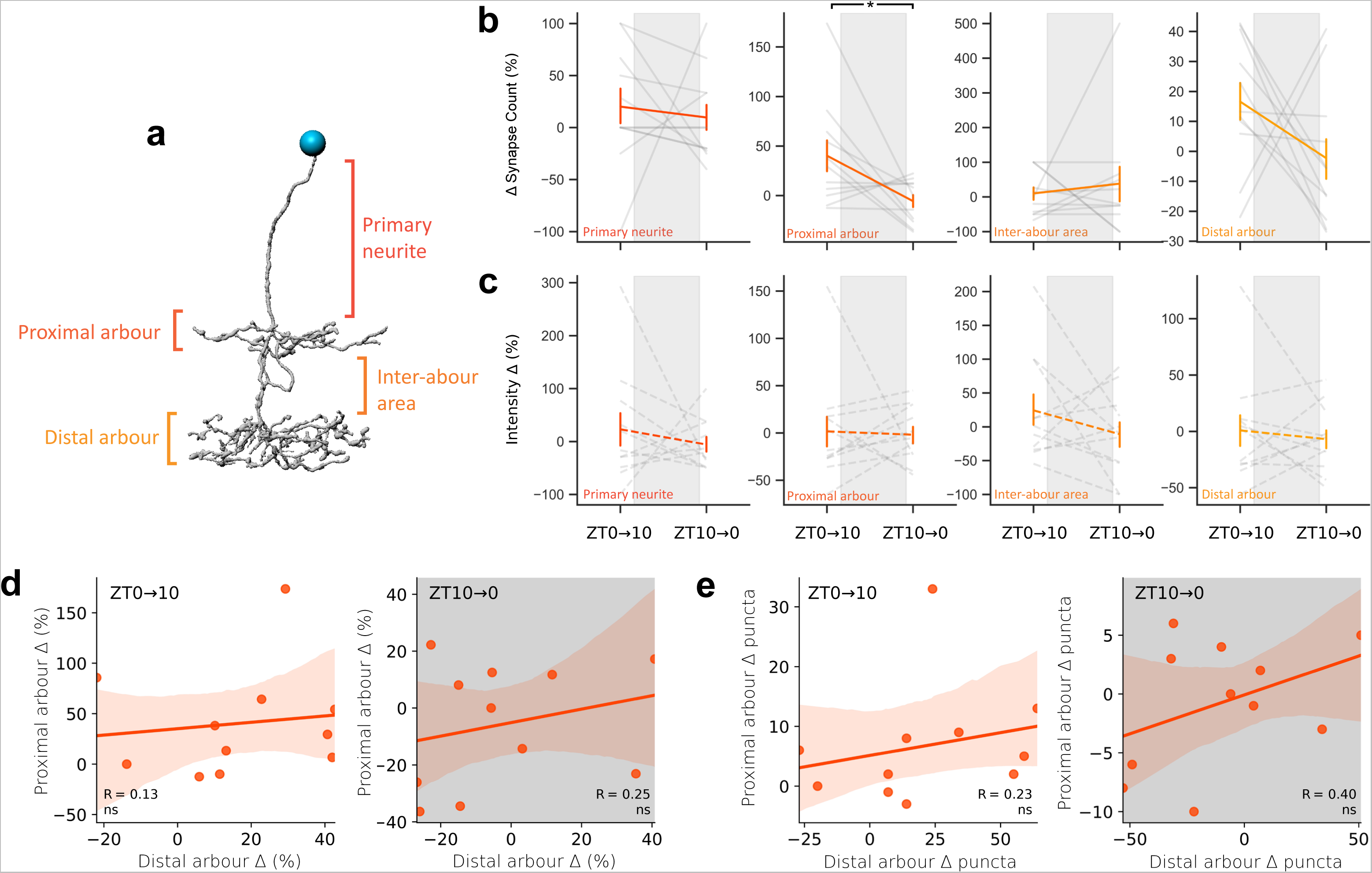
Neither sleep/wake states nor time of day have uniform effects on synapse dynamics within neuron compartments. **a**, Type 2 tectal neurons were divided into four segments: the primary neurite, proximal arbour, inter-arbour area, and distal arbour. **b-c**, The average and 68% CI of synapse number and intensity dynamics within each of the four segments. Grey lines represent segments from individual neurons. *P<0.05, repeated-measures ANOVA with Greenhouse-Geisser correction. **d-e**, Proximal and distal arbours synapse number dynamics are not correlated. **d**, The relationship between the synapse number change (%) of the proximal and distal arbours of individual Type 2 neurons during the day and night phase. **e**, The relationship between absolute synapse count changes of the proximal and distal arbours of individual Type 2 neurons during the day and night phase.

**Extended Data Figure 8:**
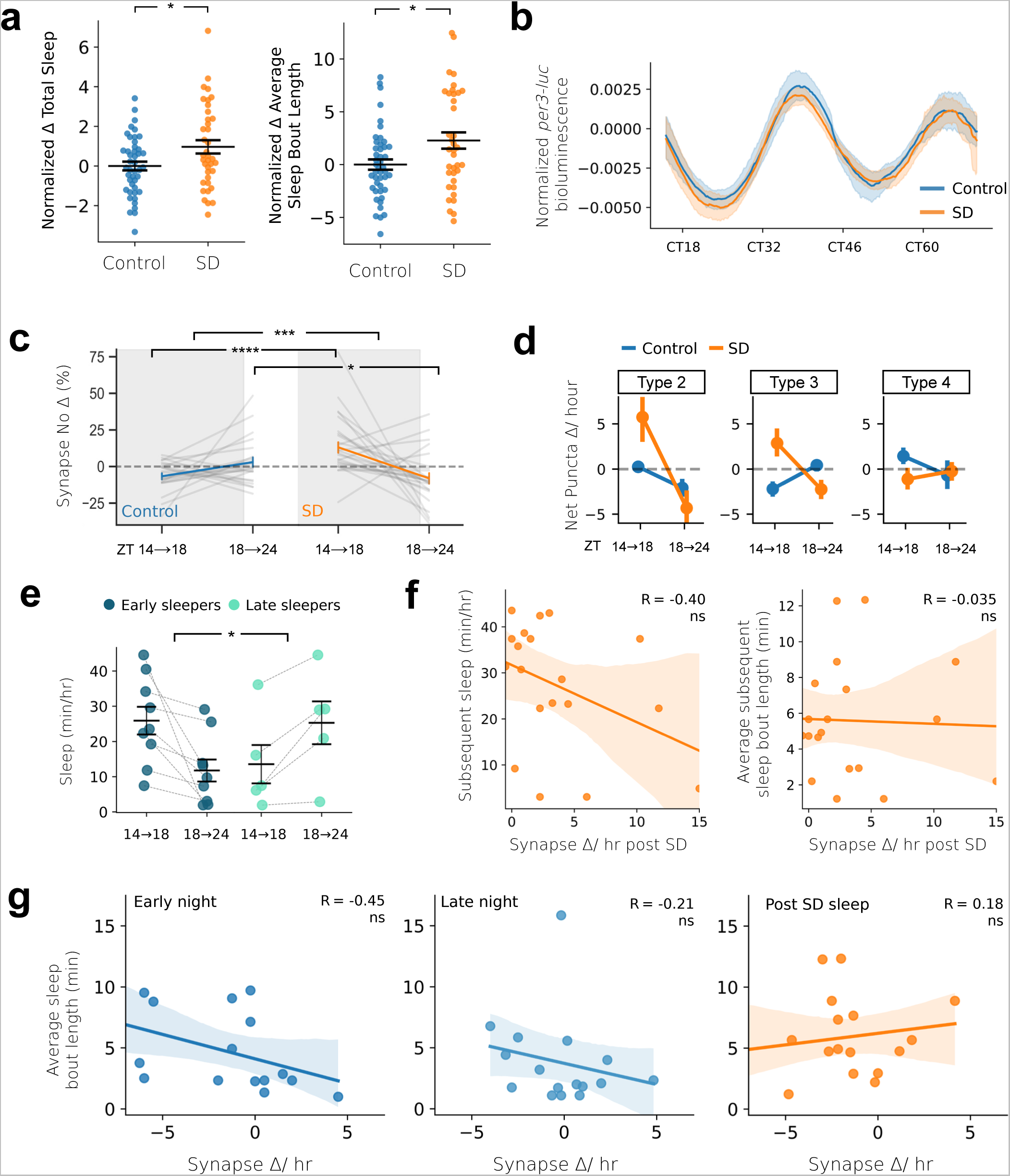
Sleep deprivation affects synapse dynamics in tectal neuron subtypes. **a**, The percentage change of total sleep (left) and average sleep bout length (right) of each larva (dots) in the 6hr post SD (ZT18-24) at 7 dpf, normalized to the circadian-matched time at 6 dpf. The black lines depict the population average ± SEM. *P<0.05, one-way ANOVA. **b**, The SD method did not alter the phase of endogenous circadian rhythms as measured by the bioluminescence driven by a *Tg(per3-luc*) reporter line for the expression of the zebrafish circadian clock gene, *per3*. The detrended *per3* bioluminescence rhythms (±95% CI) remained in phase for both SD and control larvae over multiple days of constant dark conditions. Circadian time (CT=0 last lights ON transition). **c**, The percentage change in synapse number within each neuron between imaging sessions at ZT14 and ZT18, and between imaging at ZT18 and ZT24. **d**, The average and 68% CI for net synapse change per hour for FoxP2.A tectal subtypes in control or sleep deprived larvae. **e**, Sleep amount for early and late sleepers in the early (ZT14-18) and late (ZT18-24) phase of the night. *P<0.05, ***P<0.001, ****P<0.0001, Mixed ANOVA interaction and post-hoc pairwise t-test. **f**, For each neuron/larva, changes in synapse number during extended wakefulness did not correlate with either the subsequent total sleep or average sleep bout lengths (mean ± 95% CI). **g**, For each neuron/larva, changes in synapse numbers did not significantly correlate with the average sleep bout lengths during the early and late night of controls, or after SD (mean ± 95% CI).

**Extended Data Figure 9:**
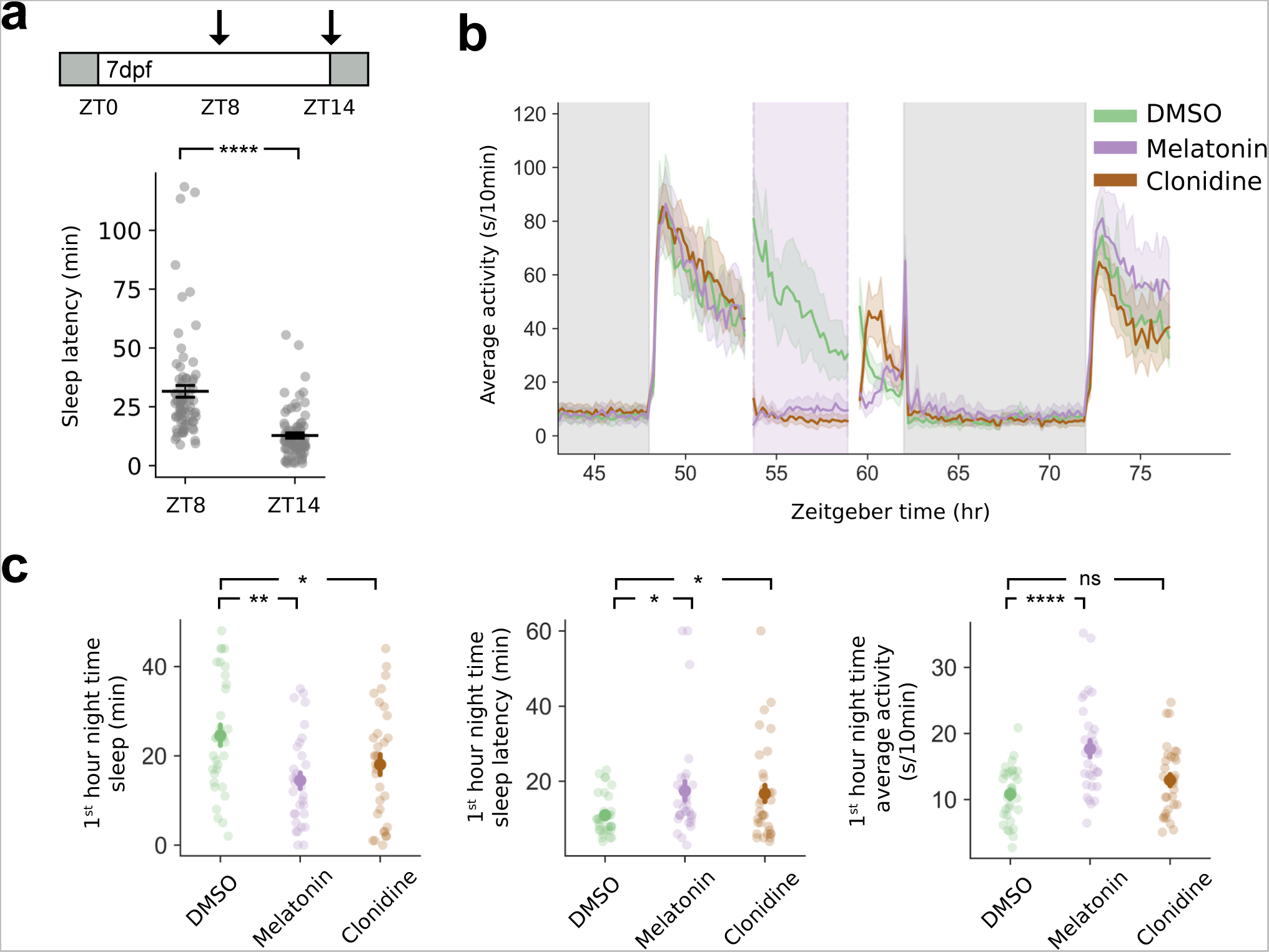
Clonidine and melatonin treatment during the day results in reduced and delayed sleep the following night. **a**, Larvae exposed to lights OFF at mid-day (ZT8, first arrow in schematic) took longer to sleep compared to lights OFF at the end of day (ZT14, 2nd arrow). ****P<0.0001, Kruskal-Wallis. **b**, Average locomotor activity on a 14hr:10hr LD cycle before, during, and after a 5hr midday (ZT5-10, 7 dpf) exposure to melatonin, clonidine, or DMSO (shaded purple panel). **c**, Larvae treated with either melatonin or clonidine from ZT5-10 had reduced and delayed sleep in first hour of the night (ZT14-15) compared to controls. *P<0.05, **P<0.01, ****P<0.0001 Dunnett’s Test.

**Extended Data Figure 10:**
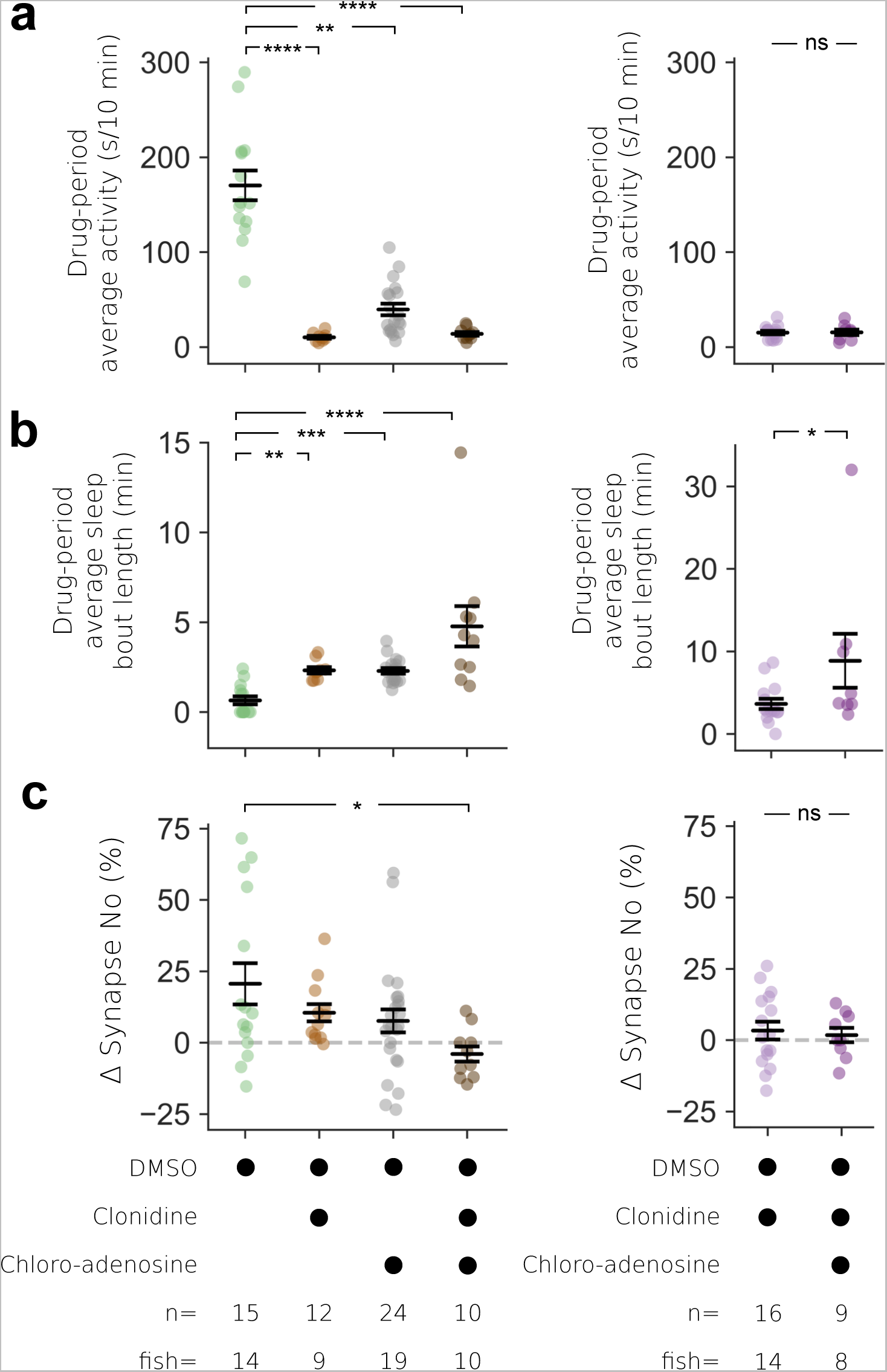
Drug-evoked day time sleep induces synapse loss only when clonidine and 2-chloroadenosine are co-administered. **a-b**, Clonidine-, 2-chloroadenosine-, and/or melatonin-treated larvae have a lower average activity and longer average sleep bout lengths during the 5hr drug period compared to DMSO treated controls. **c**, The average percentage change in synapse number (± SEM) within each neuron of DMSO, clonidine-, 2-chloroadenosine-, and/or melatonin-treated larvae. *P<0.05, **P<0.01, ****P<0.0001 Kruskal-Wallis with post-hoc Dunn’s test.

**Extended Data Figure 11:**
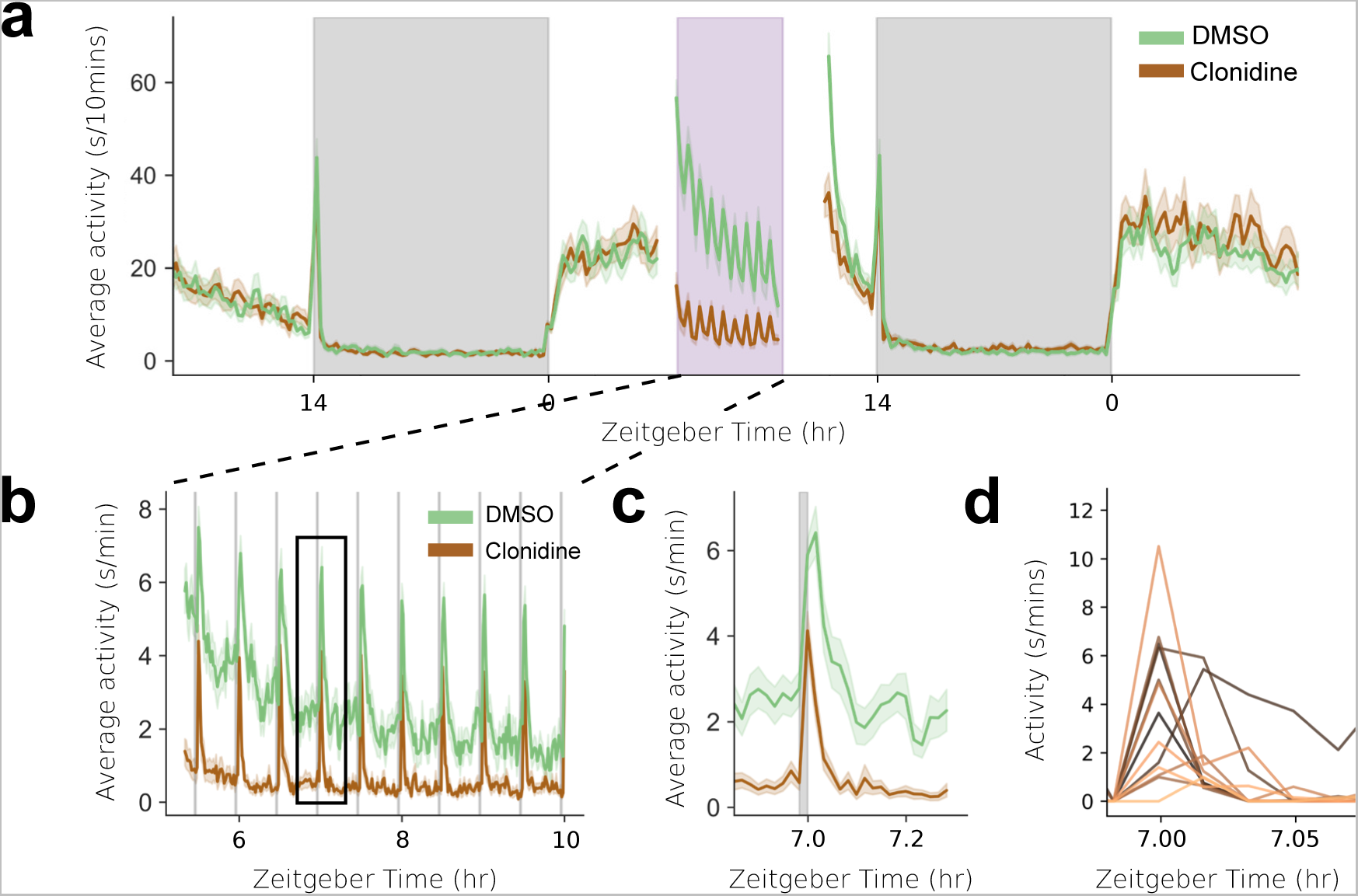
Clonidine-treated larvae respond strongly and rapidly to dark flashes. **a**, The average activity of larvae before, during and after treatment with either 30 µM clonidine or DMSO from ZT5-10 (purple shaded area) at 7 dpf. 1-minute dark pulses were given every 30 minutes during the treatment period to test for responsiveness. **b**, Higher resolution time-course of average locomotor activity during the drug treatment and dark-pulse period (ZT5-10). **c**, Both clonidine and DMSO-treated larvae respond to dark pulse with an increase in locomotion, known as the visuomotor response or dark photokinesis. Shown is the average locomotor response to a single 1-minute dark pulse delivered at ZT7. **d**, The locomotor activity for each larva-treated with clonidine (1-minute bin) at the time of dark pulse (ZT7) shown in **c**. Of the 13 larvae that were inactive at the onset of the 1-minute dark pulse, 12 rapidly increased their locomotor activity within 1 minute.

**Extended Data Figure 12:**
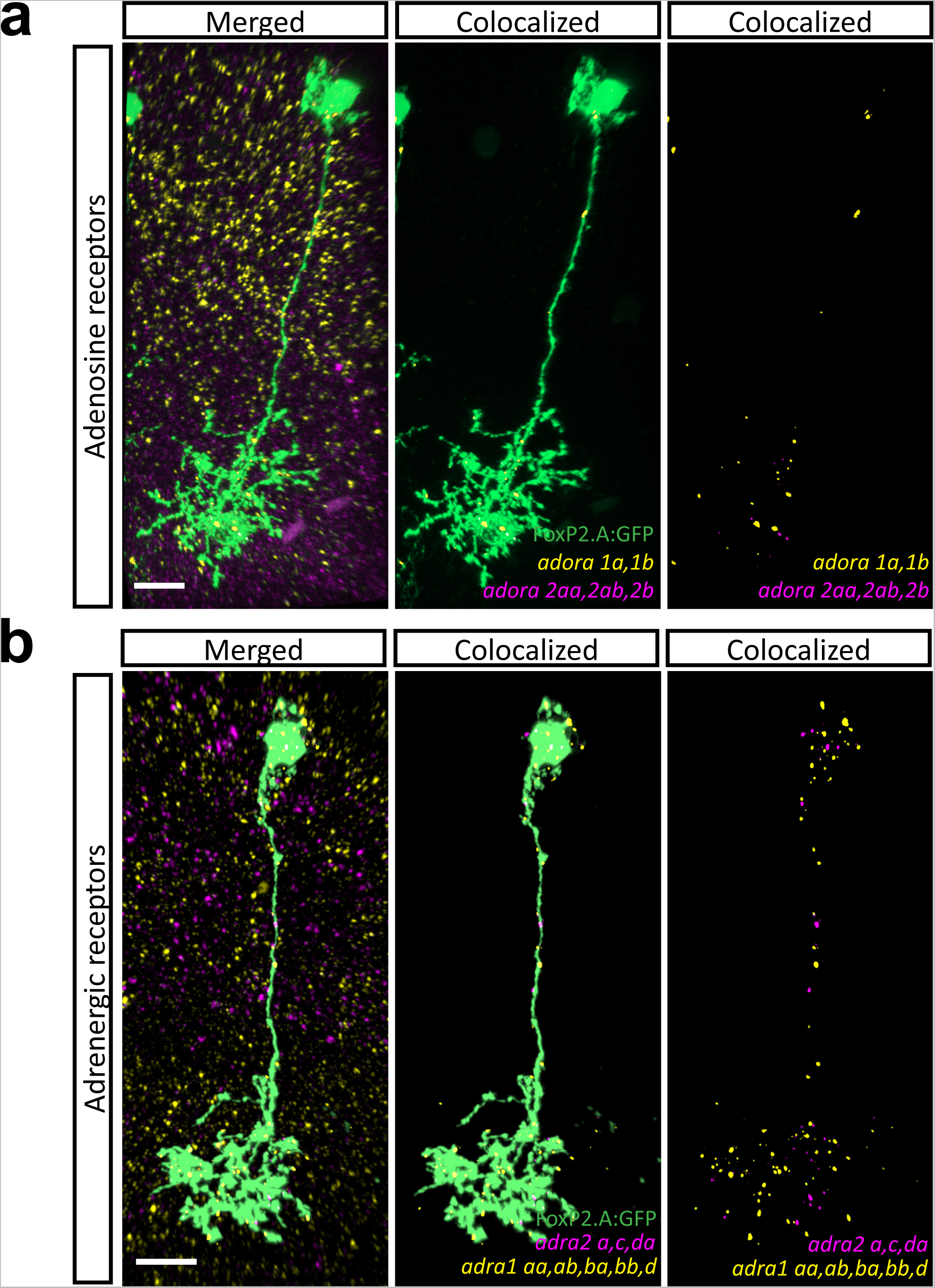
FoxP2.A+ neurons express adenosine and adrenergic receptors transcripts. Examples of adrenergic and adenosine receptor transcripts that colocalize with labelled FoxP2.A+ neurons (middle and right panel) as detected by *in situ* Hybridization Chain Reaction (HCR, see **Methods**). **a**, A single labelled tectal neuron (green) colocalizes with a cocktail of HCR probes that detect *adora1a-b* (yellow, encoding for adenosine receptors A1a and A1b) and *adora2aa*, *-ab*, *-b* (magenta, encoding for adenosine receptors A2aa, A2ab, and A2b) transcripts. **b**, Single FoxP2.A+ neuron (green) also colocalize with an HCR probe cocktail that detects *adra1 aa*,*-ab*, *-ba*, *-bb*, *-d* (yellow, encoding zebrafish α1 adrenergic receptor orthologs) and *adra2a*, *-c*, *-da* (magenta, encoding zebrafish α2 adrenergic receptor orthologs) transcripts. Scale bar, 10μm.

**Extended Data Video 1: Example of gentle handling SD.** Larvae in individual wells were manually kept awake with a paintbrush for 4 hours under red light at the beginning of the night (ZT14-18, see **Methods**). Note that many, if not most, interventions did not require physically touching the animal.

**Extended Data Table 1:**
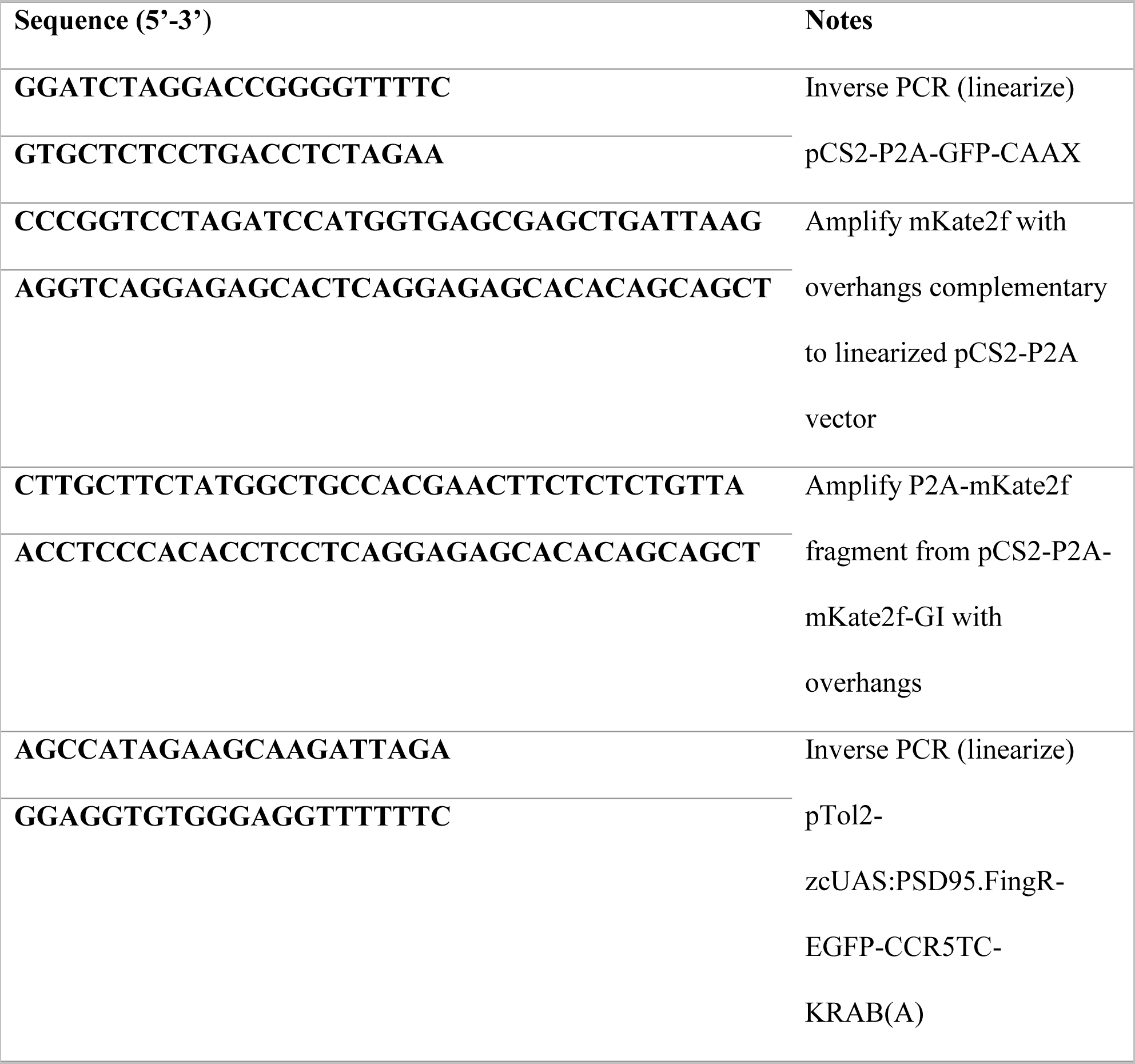
Primers used in generating DNA construct using the In-Fusion Kit.

**Extended Data Table 2:** Light schedules and imaging times in different experiments.

| **Experiment** | **Light:Dark schedule (hrs)** | **Imaging ZT and dpf** |
| --- | --- | --- |
| **Day/Night** | 14:10 (LD), 24:0 (LL),<br>LD→LL at 6 dpf (FR) | ZT0-2, ZT10-12; 7-9 dpf |
| **Overnight** | 14:10 | ZT13-14, ZT18-19, ZT0-1; 7<br>dpf |
| **Daytime drug**<br>**treatment** | 14:10 | ZT4-5, ZT10-11; 7 dpf |

